# Fusion crystallization reveals the behavior of both the 1TEL crystallization chaperone and the TNK1 UBA domain

**DOI:** 10.1101/2023.06.14.544429

**Authors:** Supeshala Nawarathnage, Yi Jie Tseng, Sara Soleimani, Tobin Smith, Maria J Pedroza Romo, Wisdom Oshireku Abiodun, Christina M. Egbert, Deshan Madhusanka, Derick Bunn, Bridger Woods, Evan Tsubaki, Cameron Stewart, Seth Brown, Tzanko Doukov, Joshua L. Andersen, James D. Moody

## Abstract

Human thirty-eight-negative kinase-1 (TNK1) is implicated in cancer progression. The TNK1-UBA domain binds polyubiquitin and plays a regulatory role in TNK1 activity and stability. Sequence analysis suggests an unusual architecture for the TNK1 UBA domain, but an experimentally-validated molecular structure is undetermined. To gain insight into TNK1 regulation, we fused the UBA domain to the 1TEL crystallization chaperone and obtained crystals diffracting as far as 1.53 Å. A 1TEL search model enabled solution of the X-ray phases. GG and GSGG linkers allowed the UBA to reproducibly find a productive binding mode against its host 1TEL polymer and to crystallize at protein concentrations as low as 0.1 mg/mL. Our studies support a mechanism of TELSAM fusion crystallization and show that TELSAM fusion crystals require fewer crystal contacts than traditional protein crystals. Modeling and experimental validation suggest the UBA domain may be selective for both the length and linkages of polyubiquitin chains.

## Introduction

Human thirty-eight-negative kinase-1 (TNK1) is implicated in cancer progression. TNK1 was discovered decades ago in human umbilical cord hematopoietic stem cells but is also expressed in other tissues, including bone marrow, and adult blood sub-populations^1^. The function of TNK1 is still poorly understood. A constitutively active, truncated TNK1 was identified in a Hodgkin’s Lymphoma cell line, driving proliferation and survival in these cells^2, 3^. Genetic screens identified TNK1 as a driver of cell survival during chemotherapy in pancreatic cancer^4^ and multiple myeloma cell lines^5^. In addition, a variety of primary human hematological cancers were shown to be dependent on TNK1 for growth^6^. In contrast, whole body deletion of TNK1 in mice results in spontaneous tumor formation^7^, whereas other in vitro studies and disease models suggest a role for TNK1 in inflammatory signaling^8–11^.

Based on sequence homology, TNK1 is grouped into the ACK family of mammalian non-receptor tyrosine kinases (NRTKs), which in humans only includes TNK1 and ACK1. Both kinases share a putative C-terminal ubiquitin-association (UBA) domain, an unusual feature among kinases^12^. UBA domains interact noncovalently with polyubiquitin and are more commonly found on de-ubiquitinases and E3 ubiquitin ligases, where they often tether the enzyme to polyubiquitin scaffolds and substrates^13^. Some UBA domains show preference for particular polyubiquitin linkages (e.g., K63-linked chains), whereas others interact indiscriminately with all known ubiquitin linkages^13–16^. Interestingly, mutations within the UBA domain of ACK1 resulted in impaired EGFR downregulation and sustained activation of downstream signaling, characteristic of an oncogenic driver^17^. In our recent study, we found that loss of an inhibitory 14-3-3 docking motif immediately adjacent to the UBA domain hyperactivates TNK1, whereas deletion of the UBA impairs TNK1 activity and alters its phospho-substrate networks^6^. These data suggest that the interaction of TNK1 with polyubiquitin is essential for normal TNK1 function. Furthermore, we found that the TNK1-UBA domain has a high affinity for multiple ubiquitin linkages and, based on sequence prediction, may have an unusual 5-helix structure^6^. In this study, we aimed to gain detailed understanding of the TNK1-UBA domain structure and the nature of its interactions with polyubiquitin. As the 9 kDa UBA domain is too small for high-resolution structure determination by cryo-electron microscopy, we used X-ray crystallography to determine its structure.

X-ray diffraction or micro-electron diffraction from protein crystals are powerful tools for determining the atomic-level structural information needed in modern structural biology^18^ but these techniques are critically dependent on successful protein crystallization. Obtaining diffraction-quality protein crystals often involves a laborious, lengthy, and expensive process of experimentally optimizing variables such as temperature, pH, ionic strength, protein concentration, crystallization reagent choice and concentration^19^. Frustratingly, such traditional protein crystallization methods successfully produce crystals for approximately 30% of input proteins and lead to an even smaller fraction of crystals that diffract well enough for atomic resolution structure determination^20^. Protein crystallization thus remains a bottleneck in protein structure determination, and lack of X-ray-quality protein crystals impedes the structure-function studies of many proteins^21–23^.

In addition to seeking the atomic structure of the TNK1-UBA domain, we sought to further our understanding of the usefulness and best use practices of TELSAM as a protein crystallization chaperone. We previously showed that 1TEL, which displays six copies of a target protein per turn of the helical axis of the TELSAM polymer, accelerated the crystallization of a fused target protein in the absence of direct inter-polymer contacts and at protein concentrations as low as 1 mg/mL^24^. Rather than solely providing new crystal contacts, TELSAM orders many copies of the target protein along and around the protein polymer, pre-programming the symmetry of the resulting crystal lattice and conferring substantial avidity to strengthen subsequent weak crystal contacts made by the target protein. TELSAM thus reduces the entropic cost of crystal nucleation and growth by pre-freezing some of the degrees of freedom. We set out to determine whether fusion to 1TEL could be effective for other proteins of interest, the lower protein concentration limit for 1TEL–target protein crystallization, and whether the flexible 1TEL–target protein linker could be extended without impairing crystal quality.

## Results

An engineered 1TEL-GG-UBA construct readily crystallized. We generated a homology model of the UBA domain and modeled ever closer superpositions of the 1TEL C-terminal and UBA domain N-terminal α-helices until the 1TEL and UBA domains were as close as possible without serious clashes between them (**Figure 1A**). This superposition left two helical residues between the C-terminal Ile-Leu of the 1TEL and the N-terminal Glu-Leu-Gln of the UBA model, which were changed to glycines to produce a flexible linker. The construct was cloned and the protein was produced and crystallized at 15 mg/mL as described above. Tapered hexagonal prism crystals of 1TEL-GG-UBA appeared in three days in four distinct crystallization conditions (**Figure 1B**). A PAGE gel of washed crystals indicated the entire construct was intact. Crystals of 1TEL-GG-UBA diffracted to an average resolution of 2.2 Å (1.78–2.54 Å across 10 crystals), indexed an average of 56% of reflections (19– 89%), had an average mosaicity of 0.66° (0.18–1.50°), and had an average data processing Isa of 20 (11–30) (**Table S1–2**).

**Figure 1:**
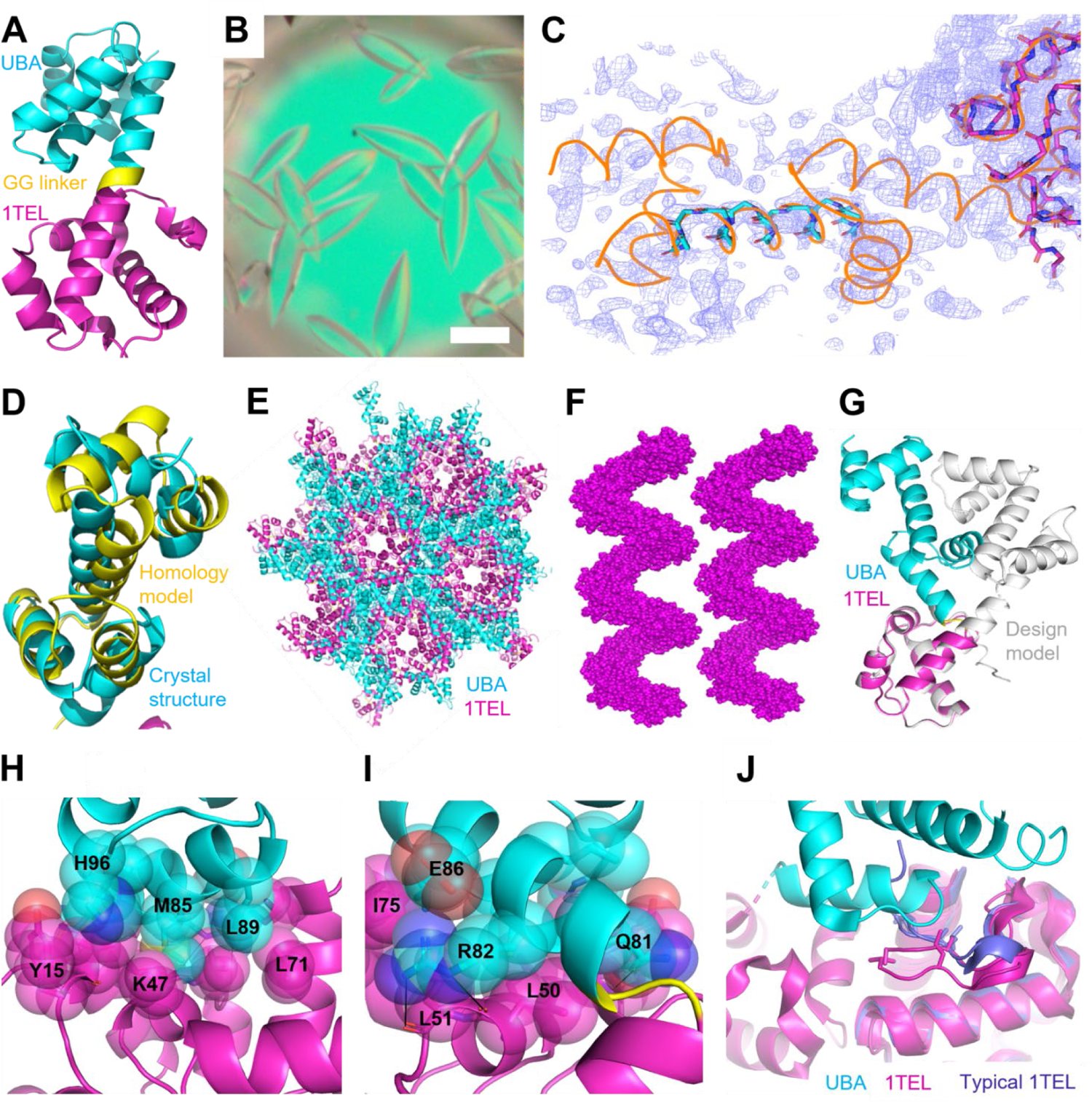
Design, crystallization, and structure of 1TEL-GG-UBA. **A.** Design model of 1TEL-GG-UBA. **B.** Representative crystals of 1TEL-GG-UBA at 15mg/mL. Scale bar is 100 μm. **C.** 1TEL-GG-UBA molecular replacement solution showing the backbones of the 1TEL domain (magenta), the UBA central α-helix (cyan), the eventual position of the UBA main chain (orange ribbon), and electron density corresponding to the initial molecular replacement solution (purple mesh, contoured to 3σ). **D.** Superposition of the homology model (yellow) onto the UBA domain (cyan) from the 1TEL-GG-UBA crystal structure. **E.** View of the 1TEL-GG-UBA crystal lattice parallel to the helical axis of the TELSAM polymers, colored as in A. **F.** View of two 1TEL-GG-UBA polymers perpendicular to their helical axes, in space filling representation and omitting the UBA domains for clarity. **G.** Superposition of the 1TEL-GG-UBA crystal structure (magenta and cyan) onto the design model (white), aligned through only the 1TEL domains. **H-I.** Two views, 180° apart, of the binding interface between the UBA domain (cyan) and the 1TEL polymer (magenta). Hydrogen bonds are shown as black lines while atoms making van der Waals contacts are shown as spheres. **J.** Superposition of the 1TEL domains of representative 1TEL wwPDB structures (purple, PDB IDs 8FT8, 8FT6^25^) onto structures of 1TEL-GG-UBA (magenta and cyan). The two 1TEL domains from 1TEL-GG-UBA are shown (magenta), while only a single UBA domain is shown (cyan). Selected amino acid side chains are shown as sticks.

TELSAM can serve as a molecular replacement search model for sufficiently small target proteins. We attempted to solve the 1TEL-GG-UBA structure using molecular replacement by placing one copy of the known 1TEL structure (PDBID: 2QAR) and 1 copy of a TNK1-UBA homology model. Phaser readily placed one copy of the 1TEL structure but was unable to locate the UBA homology model. We next tried individually placing the five α-helices of the UBA model (along with the 1TEL structure) and were able to locate the long central (3^rd^) α-helix of the UBA model, allowing us to iteratively build the remaining four UBA α-helices one at a time into the electron density map, followed by the loops between each pair of α-helices (**Figure 1C**). The resulting UBA structure was used as a molecular replacement search model for a higher-resolution dataset of the UBA crystallized on its own (described in detail below). Refinement of this later UBA-alone structure revealed that the central α-helix of the UBA from the initial 1TEL-GG-UBA structure was incorrectly displaced around it’s α-helical axis by ∼100° (1 amino acid). This central UBA α-helix was then corrected in the initial 1TEL-GG-UBA structure. The homology model created using an online server (Robetta, pre-TR-Rosetta)^26^ differed from the solved structure of the UBA domain with a Cα RMSD of 5.261 Å and was thus not accurate enough to perform successful molecular replacement. (**Figure 1D**).

TELSAM polymers do not make direct inter-polymer crystal contacts in 1TEL-GG-UBA crystals, a feature previously seen in other 1TEL-fusion crystals^24^ (**Figure 1E–F**). This result confirms that TELSAM is capable of crystallizing proteins too large to permit direct inter-TELSAM contacts. In the 1TEL-GG-UBA crystal structure, the flexibly fused UBA adopts a stable binding mode against the TELSAM polymer, also as previously seen^24^ (**Figure 1G–I**). The UBA:1TEL interface buries 1890 Å^2^ of solvent accessible surface area (average of both sides of the interface) and is mostly hydrophobic, involving residues R82, M85, L89, and H96 on the UBA and I14, L15, W27, L51, K47, L71, and I75 on the 1TEL. This observation confirms that flexibly-fused target proteins can become rigidly positioned through binding to their host TELSAM polymers and that direct inter-TELSAM contacts are dispensable if a rigid transform is maintained between adjacent TELSAM polymers. Surprisingly, this 1TEL:UBA binding mode displaces the N-terminal loop of 1TEL from the position seen in the majority of 1TEL crystal structures (**Figure 1J**), suggesting that purification tag removal prior to crystallization may be necessary for at least some TELSAM fusion constructs.

1TEL-GG-UBA can crystallize at very low protein concentrations. We previously observed that the vWa domain of human capillary morphogenesis gene 2 could be crystallized via TELSAM fusion at concentrations as low as 1 mg/mL^24^. We sought to determine whether crystallization of the TNK1-UBA domain could also occur at low protein concentrations and to probe the lower limit of necessary protein concentration. Crystals of the UBA domain on its own were seen at protein concentrations of 3.1 mg/mL. When crystallization trials of 1TEL-GG-UBA were carried out at protein concentrations of 0.1, 0.2, 0.5, 1, 2, 5, 10, and 15 mg/mL, protein crystals readily formed at all protein concentrations within three days of crystal tray setup, in Bis-Tris Mg-formate optimization trays. We selected larger crystals from many of these concentrations for X-ray diffraction. Crystals of 1TEL-GG-UBA obtained at lower protein concentrations had a similar crystal morphology to those obtained at 15 mg/mL (**Figure 2A–B**) and similar crystal quality statistics (**Table S2**). We were able to mount and diffract crystals obtained at protein concentrations as low as 0.2 mg/mL. Crystals obtained at 0.1 mg/mL were too small to be mounted for traditional X-ray diffraction experiments in our hands. The pH values at which crystals appeared generally decreased as the protein concentration decreased below 2 mg/mL (**Figure 2C**). At decreasing pH values, we hypothesize that the V112E polymerization trigger would become increasingly protonated, leading to a greater 1TEL subunit-subunit association rate and thus an increased polymer growth rate. Taken together, these observations suggest that either an increasing polymerization rate and/or longer polymers are needed to nucleate 1TEL-GG-UBA crystals as the protein concentration decreases below 2 mg/mL.

**Figure 2:**
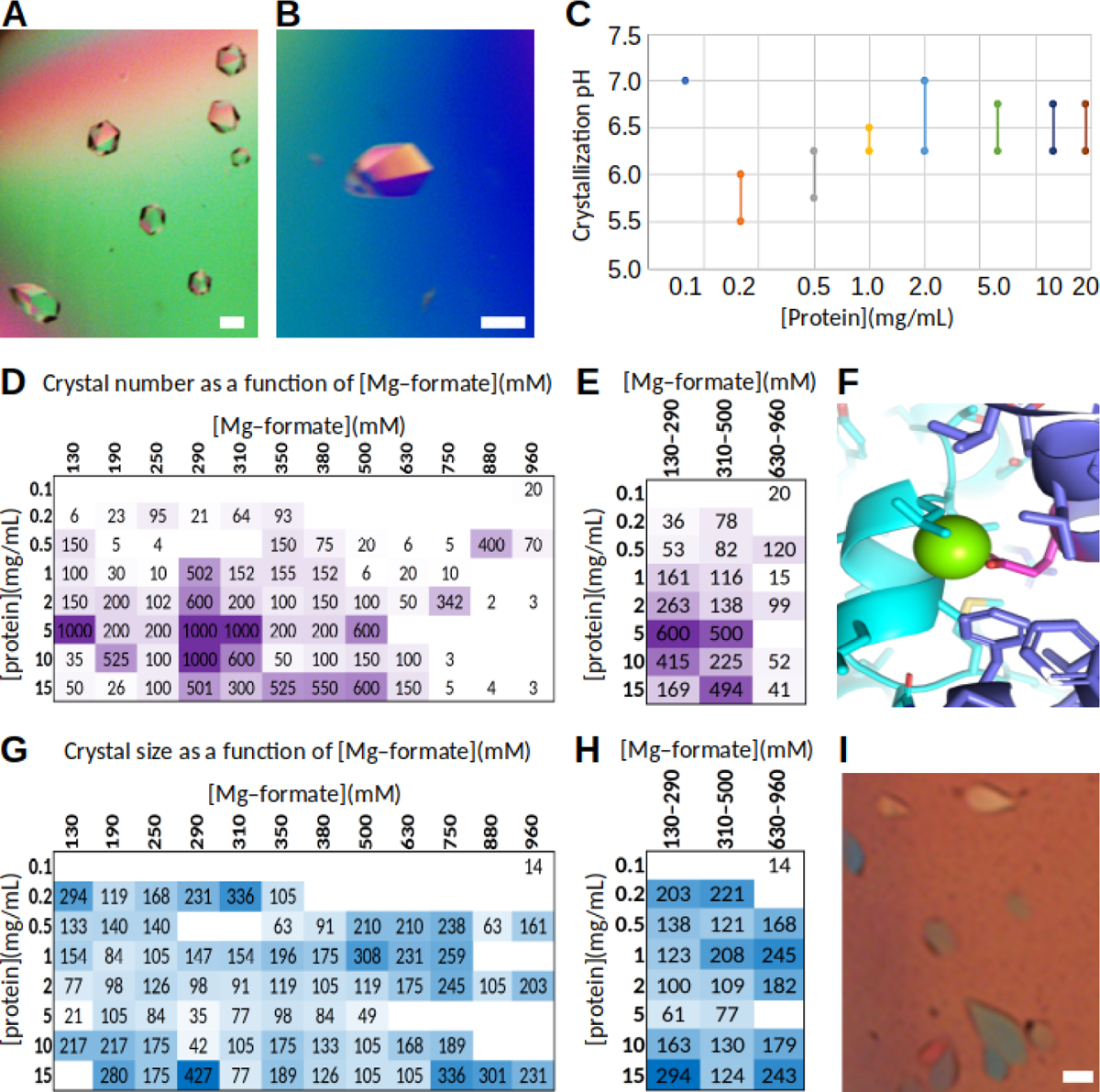
TELSAM fusion allows crystallization at low protein concentrations. **A.** Representative crystals of 1TEL-GG-UBA at 2 mg/mL. Scale bar is 100 μm. **B.** Representative crystals of 1TEL-GG-UBA at 0.5 mg/mL. Scale bar is 100 μm. **C.** pH ranges at which crystals appeared for various input protein concentrations. **D.** Heat map of the number of protein crystals observed per well as a function of Mg-formate concentration. The number of crystals observed at different pH values have been averaged. **E.** As in C. but with the Mg-formate concentrations grouped into three ranges for clarity. **F.** Model of a magnesium ion interacting with the E112 at the hydrophobic inter-subunit interface of the TELSAM polymer. Two neighboring TELSAM monomers (cyan and purple) are shown as they would appear in a polymer. The V112E substitution is highlighted in pink while the putative magnesium ion is represented by a green sphere. **G.** Heat map of the maximum crystal size (µm) observed per well as a function of Mg-formate concentration. The maximum is the largest crystal observed at that Mg-formate concentration across all pH values tested. **H.** As in G. but with the Mg-formate concentrations grouped into three ranges for clarity. **I.** Representative crystals of 1TEL-GSGG-UBA. Scale bar is 100 µm.

Mg-formate appears to inhibit 1TEL-GG-UBA crystal nucleation. An analysis of 0.1, 0.2, 0.5, 1, 2, 5, 10, and 15 mg/mL 1TEL-GG-UBA crystal trays showed a generally negative correlation between the Mg-formate concentration and the number of protein crystals per well. The reduction in crystal number by increasing Mg-formate concentration was most evident at Mg-formate concentrations > 310 mM and protein concentrations > 0.5 mg/mL (**Figure 2D–E**). This result contrasts with the behavior of traditional protein crystallography precipitants, which often result in greater numbers of crystals at higher precipitant concentrations. We suggest that a concentration-dependent transitory interaction between Mg^2+^ ions and the negatively-charged sidechain carboxylate of the crucial 1TEL V112E substitution may occur. This E112 lies at the hydrophobic inter-subunit interface and TELSAM polymerization can theoretically only occur when it becomes protonated and neutrally-charged at lower pH values^27^. The hydrophobic nature of the TELSAM inter-subunit interface would favor interaction between the Mg^2+^ ions and the carboxylate group. The interaction between the carboxylate and Mg^2+^ would sterically block TELSAM subunit association and thereby result in decreased rates of TELSAM polymerization and crystal nucleation (**Figure 2F**). This steric hinderance would be greater if the Mg^2+^ ions were in a hydrated state when binding to the V112E carboxylate^28, 29^. We observed some degree of protein precipitation at protein concentrations ≥ 15 mg/mL and Mg-formate concentrations > 750 mM, which may also have resulted in the smaller numbers of crystals in these conditions. The crystals formed at Mg-formate concentrations as low as 130 mM are particularly noteworthy. They show that 1TEL-GG-UBA is capable of forming diffraction-quality crystals in the virtual absence of precipitants (because this Mg-formate concentration is within the range of the NaCl or KCl salts in typical biological buffers). This behavior was not seen when the UBA domain was crystallized on its own.

The number of 1TEL-GG-UBA crystals was largely independent of the input protein concentration until protein concentrations < 0.2 mg/mL were sampled. We also observed that the time to crystal appearance appeared to also be largely independent of protein concentration. This reveals that 1TEL-GG-UBA experiences a similar number of nucleation events per unit volume at a wide range of protein concentrations (0.2–15 mg/mL), suggesting that nucleation is not protein concentration-dependent. Based on this and the above observations, we hypothesize that 1TEL-GG-UBA crystals do not nucleate as traditional protein crystals do, by confining enough copies of the protein into a small space to yield a protein-protein association rate sufficient to form a stable crystal nucleus. This is consistent with the avidity conferred to the UBA domains (via fusion to 1TEL polymers) playing a significant role in crystal nucleation. We propose that 1TEL-GG-UBA polymers zipper up in response to the avidity of the UBA domains. Surprisingly, we also did not observe any clear trends in crystal size as a function of Mg-formate concentration (**Figure 2G–H**).

Limited extension of the 1TEL–UBA linker can be tolerated in 1TEL–UBA crystals. We were intrigued that a flexible linker did not abrogate crystallization of 1TEL-flex-vWa or 1TEL-GG-UBA. We sought to determine the maximal flexible linker length that would still allow the formation of well-ordered 1TEL–UBA crystals. We designed and cloned a series of constructs that had 4, 6, 8, or 10 amino acids between the 1TEL C-terminus and the UBA N-terminus, using glycines interspersed with serines (1TEL-GSGG-UBA, 1TEL-GSGGSG-UBA, 1TEL-GGSGGSGG-UBA, and 1TEL-GSGGSGGSGG-UBA). We produced and crystallized these proteins at a concentration of 20 mg/mL. Crystals of the 1TEL-GSGG-UBA construct appeared in 10–14 days in numerous conditions and had a similar crystal morphology to the 1TEL-GG-UBA crystals (**Figure 2I**) and similar crystal quality statistics (**Table S2**). Crystals of 1TEL-GSGGSG-UBA appeared in 85 days in a single condition, exhibited severe non-merohedral twinning, and did not diffract. Crystals were not obtained for 1TEL-GGSGGSGG-UBA or 1TEL-GSGGSGGSGG-UBA, even after an extended period. These results suggest that while the length of the flexible 1TEL–target protein linker can be extended somewhat without abrogating the formation of well-ordered crystals, there is an upper limit. The 1TEL-GSGG-UBA construct took 3–5 times as long to crystallize as the 1TEL-GG-UBA constructs, while the 1TEL-GSGGSG-UBA construct took ∼30 times as long (**Table S2**). As a longer 1TEL–target protein linker would be expected to slow the on-rate of UBA association with the host 1TEL polymer, this result suggests that UBA association with the host 1TEL polymer is likely a rate-limiting step for inter– polymer association and crystal formation.

All 1TEL–UBA crystal structures have nearly identical UBA binding modes against the 1TEL polymer. A single dataset from each of the 0.5 and 2 mg/mL 1TEL-GG-UBA crystallization conditions and from the 1TEL-GSGG-UBA construct was selected for structure solution, striking a balance between best possible resolution, dataset quality, representation of various protein concentrations, and representation of the varying TELSAM helical rise (**Figure S1**). The X-ray phases were solved by molecular replacement using one copy of the 1TEL domain (PDBID: 2QAR)^30^ and the UBA domain from the UBA-alone structure (described below). Aligning the 15 mg/mL, 2 mg/mL, and 0.5 mg/mL 1TEL-GG-UBA and the 1TEL-GSGG-UBA structures through only their 1TEL domains results in a nearly identical placement of the UBA domains, with an average UBA domain Cα RMSD of 0.967 Å (0.594 to 1.410 Å) (**Figure 3A**). Concomitant with the conserved UBA:TELSAM *cis* binding mode, the UBA:TELSAM *cis* interface packing was similar in all structures, with UBA M85 and L89 (1TEL-GG-UBA numbering) forming hydrophobic interactions with the host 1TEL I14, K47, L50, and L51 and with W27, F32, and L71 of a neighboring 1TEL monomer within the same 1TEL polymer (**Figure 3B**). Despite the fact that most of the UBA sidechain conformations in this region are highly conserved (excepting the C-terminus of the neighboring 1TEL molecule), the conformation of the UBA M85 is highly variable between the four structures and the Cε atom is only visible in the 1TEL-GSGG-UBA structure. The conformation of the UBA L89 is more conserved among the four structures, although this side chain is not visible in the 15 mg/mL structure (**Figure 3B**). These observations suggest that the UBA observed binding mode is significantly lower in energy than other potential binding modes, in contrast to the many binding modes observed for 1TEL–CMG2_vWa fusions^25^. Concomitantly, twinning, translational pseudosymmetry, and TELSAM polymer flipping were not observed in any of our 1TEL–UBA crystals.

**Figure 3:**
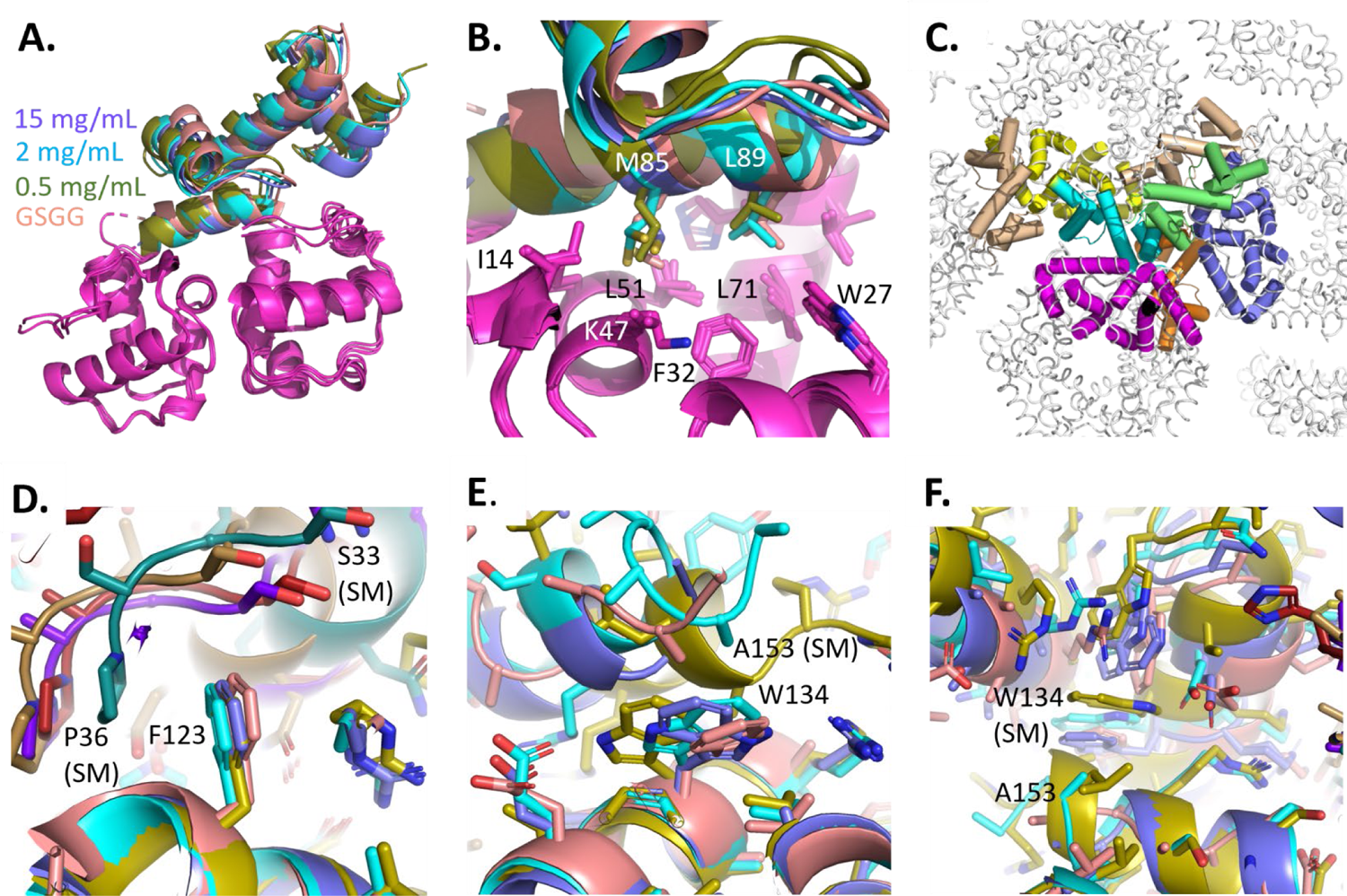
1TEL-UBA structures have nearly identical UBA binding modes against the 1TEL polymers. **A.** 15 mg/mL (purple), 2 mg/mL (cyan), and 0.5 mg/mL (olive green) 1TEL-GG-UBA structures superimposed onto the 1TEL-GSGG-UBA structure (salmon), via the 1TEL domain (magenta). **B.** As in A. but zooming in on the UBA:1TEL interface. **C.** Schematic of crystal contacts between a UBA domain (cyan), six other 1TEL subunits (yellow, purple, and magenta) and three other UBA domains (wheat, orange, and green). Domains contacting the given UBA domain are shown as cylinders and colored, while other 1TEL subunits are shown as a gray ribbon. **D. C**rystal packing differences between the four 1TEL-UBA structures, focusing on F123. The structures are colored thus: 15 mg/mL UBA (purple), 15 mg/mL 1TEL (purple), 2 mg/mL UBA (cyan), 2 mg/mL 1TEL (blue-green), 0.5 mg/mL UBA (olive green), 0.5 mg/mL 1TEL (brown), GSGG UBA (salmon), and GSGG 1TEL (firebrick). All structures have been aligned through the UBA domain that hosts the F123 shown. This same UBA domain appears in the lower portion of panels D, E, and F (SM = symmetry mate). **E.** As in D. but focusing on W134. **F.** As in D. but focusing on the C-terminus of the UBA.

The TELSAM polymers in the 1TEL-UBA crystals vary significantly in their degree of helical rise. While all of the 1TEL-UBA crystal structures have nearly identical UBA binding modes against the 1TEL polymer, the helical rise of the TELSAM polymers varies by as much as 11.6%, even differing by as much as 9.5% between crystals from a single crystallization drop. This variation in helical rise did not significantly correlate to the protein concentration (R^2^ = 0.13) nor to the diffraction limits of these crystals (R^2^ = 0.085) (**Figure S1A–C**). As previously noted, the diffraction limits of these crystals may additionally be dependent on factors such as crystal size, shape, orientation in the loop, freezing conditions, and researcher skill level.

We sought to identify the cause of this variation in 1TEL–UBA helical rise. In most 1TEL– vWa^24^ and all 1TEL-GG-UBA crystals, target proteins do not contact the next turn of their host polymer. This observation suggests that the helical rise of the TELSAM polymers is not dictated by *cis* interactions with the target protein but rather by the *trans* crystal packing between adjacent 1TEL– target protein units. Since the 1TEL polymers themselves make no direct contacts with their neighboring polymers, we analyzed instead the interactions made by the UBA domains. Each UBA domain contacts its own host 1TEL subunit, two nearby 1TEL subunits in its own host polymer, two UBA domains from its own polymer, four 1TEL subunits from two neighboring 1TEL–UBA polymers, and two UBA domains from one of these neighboring 1TEL–UBA polymers. (**Figure 3C**). Alignment of the four crystal lattices through a single UBA domain reveals subtle differences in crystal packing. For example, UBA F123 is essentially identical in all four crystal structures, but makes differing contacts with the loop containing S33 and P36 of a neighboring 1TEL domain, packing on P36 in the 2 mg/mL structure (blue-green), on S33 in the 15 mg/mL and GSGG structures (dark purple and firebrick, respectively), and on neither of these in the 0.5 mg/mL structure (brown) (**Figure 3D**). In another example, UBA W134 is seen in various side chain conformations and packs in various ways on the C-terminal α-helix of a neighboring UBA domain, which is itself positioned differently relative to the first UBA domain across the four crystal structures (**Figure 3E**). The C-termini of the aligned UBA domains across the four structures align to each other reasonably well (**Figure 3F**), indicating that the differential crystal packing of this region across the four structures is likely not a result of differential positioning of the UBA C-terminus. This analysis does not reveal whether the differential crystal packing causes the differential helical rise or vice versa, but it does demonstrate that the differential helical rise correlates with variability in the crystal contacts made by the UBA domains and does not simply represent variable expansion of the unit cell along its c-axis with no change in the crystal contacts. Curiously, the number of amino acids visible at the UBA C-terminus is strongly correlated to the input protein concentration, with the 0.5 mg/mL structure resolving all amino acids, the 2 mg/mL structure missing one amino acid, the 15 mg/mL structure missing four amino acids, and the 20 mg GSGG structure missing six amino acids. The number of resolvable amino acids at the UBA C-terminus did not correlate to the resolution of each structure, as might be expected (**Table S2**). Unlike the stronger correlation between helical rise and resolution in 1TEL–CMG2_vWa crystals^25^, the correlation was much weaker in 1TEL–UBA crystals and had the opposite trend, with larger helical rises generally giving somewhat higher resolution (**Figure S1C**). This suggests the helical rise yielding more uniform and thus better diffracting crystals is unique from construct to construct and is likely that degree of helical rise that enables the strongest-possible contacts to be made.

The variable helical rise further hints at a mechanism for the crystallization of 1TEL-UBA fusions. The fact that a single 1TEL-GG-UBA protein construct with nearly identical UBA *cis*-binding modes against the host polymer can result in polymers with significantly different TELSAM polymer helical rise and subtly different *trans* contacts between neighboring 1TEL–UBA polymers hints at flexibility in the way these polymers associate during initial crystal nucleation. We propose that the initial crystal nuclei must have been formed by 1TEL–UBA polymers that first polymerized and then zippered together via their UBA domains, even if these polymers may have been short. This initial crystal nucleus would have thus defined the specific inter-polymer crystal contacts and the degree of helical rise, which was then propagated throughout the rest of the crystal. In this model, even separate nucleation events in single drop could adopt slightly different inter-polymer contacts and helical rise. This observation also suggests that the avidity conferred to the UBA domain (through fusion to the 1TEL polymer) was sufficient to stabilize a variety of nonidentical inter-polymer crystal contacts. The energy well for 1TEL-GG-UBA polymer–polymer association has several equally low energy minima that subtly differ in their precise structures. This observation also suggests that having a reproducible target protein binding mode against the TELSAM polymer doesn’t enforce a reproducible polymer– polymer orientation. If it is possible to have subtly different *trans* polymer–polymer contacts between different crystals of the same construct, it is conceivably possible to have such subtle differences within a single crystal. While one might expect that such differences in crystal packing might abrogate crystal growth, we previously showed that TELSAM fusion can form crystals dependent on extremely weak crystal contacts^24^, having multiple target protein binding modes in the same crystal, or having multiple distinct polymer–polymer interactions in the same crystal, e.g., polymer flipping^25^. These examples suggest the possibility of obtaining diffracting crystals of TELSAM–target protein fusions that exhibit internal heterogeneity. If these internal packing differences were sufficiently large, such crystals would be expected to exhibit poorer diffraction limits or no diffraction at all. Based on the observations above, we propose that the formation of well-ordered crystals of TELSAM–target protein fusions involves three essential steps: 1. The TELSAM polymers must polymerize. 2. All fused target proteins must find a single, consistent, rigid, *cis* binding mode against their host polymers. 3. All TELSAM–target protein polymers must find a single, consistent, rigid *trans* binding mode with neighboring TELSAM–target protein polymers. The target protein most likely must choose a binding mode against its host polymer no later than the moment of polymer–polymer association. Failure in step 1 will likely result in no crystallization or at best a crystallization rate and propensity no better than those seen in traditional protein crystallography experiments. Failure in step 2 or 3 would either also result in no crystallization or could possibly result in crystals with poor resolution diffraction or no diffraction at all.

Fusion to 1TEL does not perturb the folded structure of the UBA domain. We cloned, produced, crystallized, and diffracted the UBA domain without fusion to 1TEL. Two days after setting trays, thin rectangular plate crystals of UBA-alone appeared in two conditions (**Table S2)**. This crystal habit suggests a growth defect in one dimension (**Figure 3G**) and is inferior to the bulkier, hexagonal prism crystal habit of 1TEL-UBA fusions. The UBA domain plate crystals diffracted to around 1.5 Å resolution (1.38-1.73 Å across 3 crystals, I/σ ≥ 2), exhibited low estimated mosaicity (0.3-1.3°), indexed 20-64% of reflections, and had an average data processing Isa of 19 (15–21) (**Table S2**). Molecular replacement was carried out by placing two copies of the UBA domain from the 15 mg/mL 1TEL-GG-UBA structure in to the P21212 asymmetric unit (**Figure 3H**). Following structure refinement, these two copies of the UBA domain were essentially identical to each other and to the UBA domains from the 15 mg/mL, 2mg/mL, and 0.5 mg/mL 1TEL-GG-UBA, and the 1TEL-GSGG-UBA structures (**Figure 3I**). This observation confirms that fusion to 1TEL does not perturb the structures of target proteins. We also note that these structures of the UBA domain closely match that of the AlphaFold2 prediction (Uniprot Q13470: AF-Q13470-F1), with an average Cα RMSD of 0.747 Å (0.451–0.966 Å)^31, 32^ TELSAM fusion is correlated with higher refined crystallographic B-factors in the target protein, relative to that target protein crystallized on its own. We previously observed that when fused to TELSAM, target proteins required fewer or less strong crystal contacts to form diffracting crystals.

This allowed visualization of increased protein dynamics in a DARPin target protein at its C-terminus, furthest from its connection point to the TELSAM polymer^24^. The resolution difference between the 3TEL-rigid-DARPin structure and the previously-published structure of an identical DARPin^33^ prevented direct comparison of the crystallographic B-factors between these two structures. The more similar resolution cutoffs between the UBA domain crystallized with (1.5 Å) and without (1.4 Å) fusion to 1TEL allowed us to make a more direct comparison of the degree of flexibility enabled by a TELSAM-mediated crystal lattice. The average refined B-factor for the UBA-alone crystal structure (1.4 Å) was found to be 29.35 Å^2^. Within the UBA-alone structure, B-factors appeared to be greatest in loop regions, but also in the C-terminal half of the protein (**Figure 4D**). The 2 mg/mL 1TEL-GG-UBA structure (1.5 Å) had an average refined B-factor of 40.22 Å^2^. For this 1TEL-GG-UBA structure, the B-factors were again greatest in loop regions but were significantly higher in the UBA domain (48.63 Å^2^) than in the 1TEL domain (36.41 Å^2^). B-factors were particularly high at the C-terminal region of the UBA domain, notably in residues 149–154 (68.44 Å^2^), supporting the idea that increased C-terminal B-factors are likely due to increased target protein conformational flexibility, and not due to disorder in the crystal lattice (**Figure 4E**). Analysis of the 2 mg/mL 1TEL-GG-UBA structure reveals the protruding UBA domains make fewer lattice contacts as they extend out radially from the TELSAM polymer, with a concomitant increase in B-factors (**Figure 4F**). TELSAM fusion crystallography thus appears to allow a greater degree of native protein dynamics.

**Figure 4:**
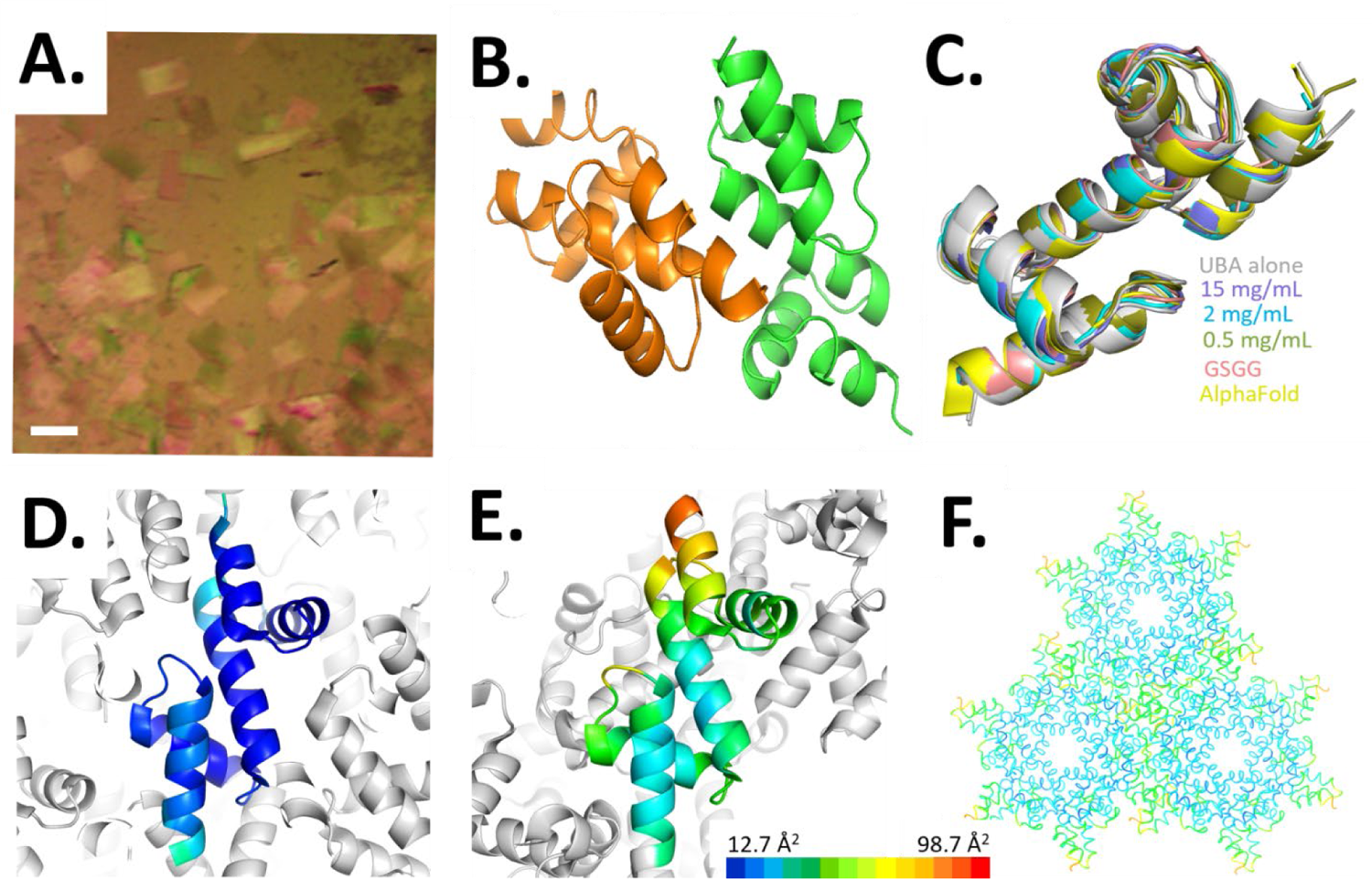
UBA-alone crystal structure and B-factor comparison. **A.** Representative crystals of the UBA domain crystallized on its own. The scale bar is 100 µm. **B.** Asymmetric unit of the UBA-alone crystal structure, with two UBA chains. **C.** Superposition of the two chains from the UBA-alone structure (white) with the UBA domains from the four 1TEL–UBA fusion structures (15 mg/mL–purple, 2 mg/mL–cyan, and 0.5 mg/mL–olive green) and the AlphaFold2 prediction–yellow. **D.** B-factors of a single chain within the UBA-alone crystal lattice, range 12.7-79.7 Å^2^. **E.** B-factors of a single chain within the 2 mg/mL 1TEL-GG-UBA crystal lattice range 27.1-86.5 Å^2^. **F.** Three 1TEL-GG-UBA polymers from the 1TEL-flex-UBA 2mg/mL crystal lattice, viewed along their helical axes and colored according to the refined B-factors. The color scale is the same in panels D-F.

TELSAM fusion crystals involve less-extensive target protein crystal contacts that crystals of those same target proteins without TELSAM. We measured the solvent-accessible surface area buried by the UBA domain in each of these structures. We ignored solvent ions and ligands and normalized the buried surface area to the number of amino acids resolved in the UBA domains from each structure. In the UBA-alone structure, interactions between the chain A UBA domain and other molecules around it buried 2828 Å^2^ of solvent accessible surface area (average of both sides of the interface), or 36.73 Å^2^ per residue, while interactions between the chain B UBA domain and other molecules around it buried 2613 Å^2^ of solvent accessible surface area, or 33.94 Å^2^ per residue. In contrast, in the 2 mg/mL 1TEL-GG-UBA structure, interactions between the UBA domain and other molecules around it (including its host 1TEL domain and linker) buried 831 Å^2^ of solvent accessible surface area, or 10.60 Å^2^ per residue. We extended this analysis to compare our recent 1TEL-TV-vWa structure (1.6 Å resolution, PDBID 8FT8^25^) with the previously published highest-resolution structure of the vWa domain crystallized on its own (1.5 Å resolution, PDB ID: 1SHU^34^. In the vWa alone structure, interactions between the vWa domain and other molecules around it buried 2074 Å^2^ of solvent accessible surface area, or 11.46 Å^2^ per residue. In the 1TEL-TV-vWa structure, interactions between the vWa domain and other molecules around it (including its host 1TEL domain and linker) buried 1678 Å^2^ of solvent accessible surface area, or 9.59 Å^2^ per residue. This is not a perfectly fair comparison because in TELSAM fusion crystals, proteins of interest are covalently tethered to their host TELSAM polymers. Indeed, this tethering of target proteins to TELSAM polymers is likely why those target proteins need less contact area per residue with other molecules in the crystal lattice. Taken together, these data reveal that TELSAM fusion allows structure determination of proteins of interest with comparable resolution to the highest resolution structures determined without TELSAM fusion, while burying 16–40% less solvent accessible surface area per residue.

1TEL-GG-UBA binds ubiquitin in vitro. We previously demonstrated that the TNK1-UBA domain can interact with various linkage variants (K48- or K63-) tetra-ubiquitin^6^. We confirmed that the UBA domain in our 1TEL-GG-UBA construct could bind to M1-linked-mono-, di-, tri- and tetra-ubiquitin using size exclusion chromatography (SEC) (Figure 5A–D). In our results, we observed 100% binding, likely because our input protein concentrations were much higher than the Kd’s of the 1TEL-GG-UBA:ubiquitin complexes^6^. We also note that the di-, tri- and tetra-ubiquitin yielded polydisperse peaks when run alone, likely due to flexible M1-linked ubiquitin adopting multiple conformational states. However, when the ubiquitin variants bound to 1TEL-GG-UBA, we observed shifts from polydisperse peaks to monodisperse peaks (Figure 5B–D). This suggests that M1-polyubiquitin adopts a single conformation upon binding to 1TEL-GG-UBA, meaning that either all four ubiquitin molecules interact with the UBA simultaneously or that a subset of them interact and the remaining non-interacting ubiquitin molecules adopt a single stable orientation relative to those that interact. The stoichiometry of UBA-polyubiquitin interaction was invariably 1:1. This observation is surprising in the case of mono and di-ubiquitin molecules, which might be expected to respectively exhibit 1:4 and 1:2 UBA:ubiquitin stoichiometries when considering that tetra-ubiquitin also has a 1:1 binding stoichiometry. This result suggests that there are not enough high-affinity ubiquitin binding sites on the UBA to allow more than one mono- or M1-di-ubiquitin to bind at a time and that there exists a single higher-affinity anchor site that a mono- or polyubiquitin chain must occupy in order to bind to the UBA.

**Figure 5:**
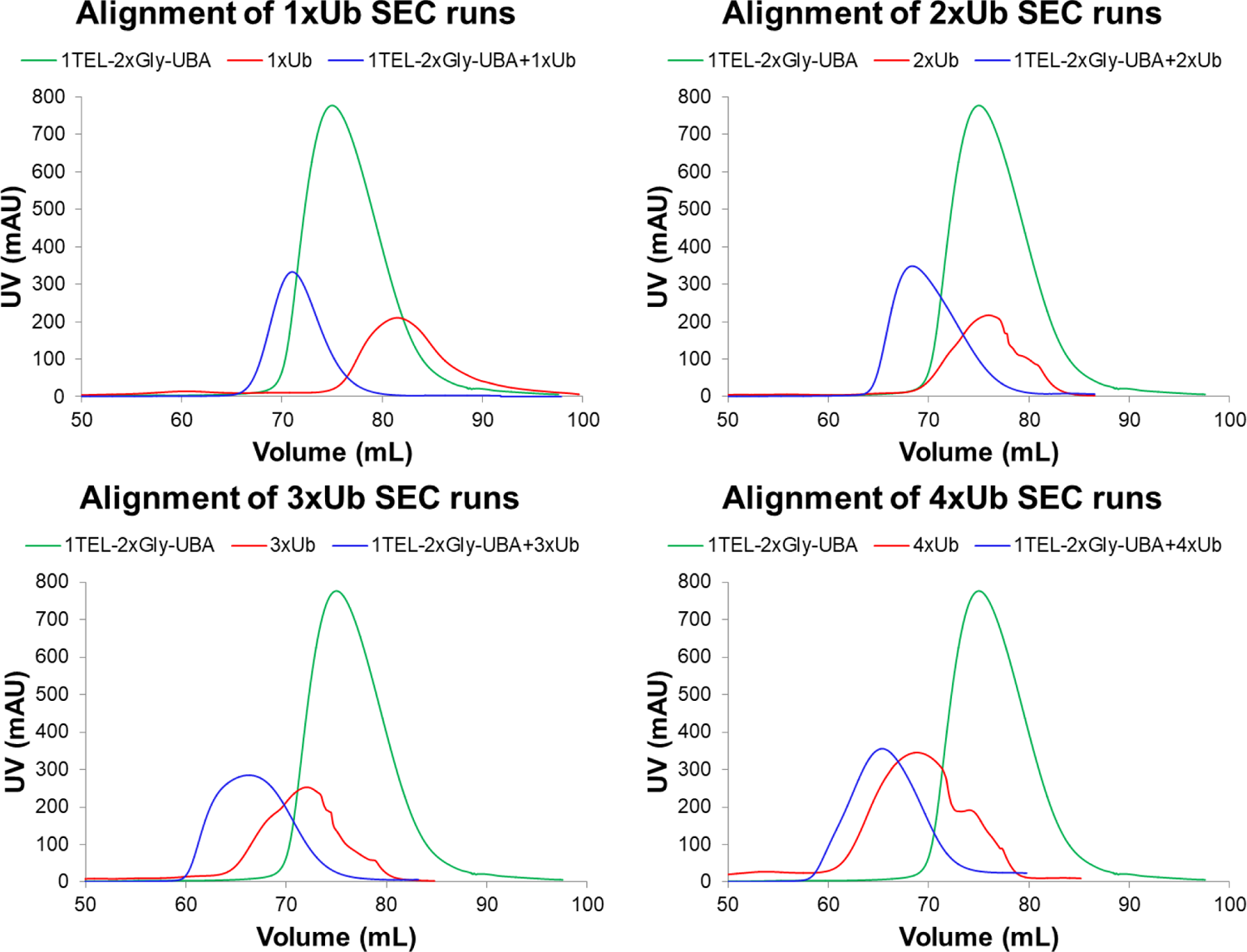
Aligned SEC runs of 1TEL-GG-UBA (green) with. **A.** mono-ubiquitin, **B.** M1-di-ubiquitin, **C.** M1-tri-ubiquitin and **D.** M1-tetra-ubiquitin with 1TEL-GG-UBA-alone (green), ubiquitin-alone (red) and combined (blue).

Modeling suggests that the TNK1-UBA domain may be selective for polyubiquitin chains. We next used the UBA-alone crystal structure to model potential binding interactions between a single ubiquitin module and the UBA domain. Fifteen UBA-Ubiquitin crystal and NMR structures from the Worldwide Protein Data Bank (wwPDB, PDBIDs: 1OTR, 1P3Q, 1WR1, 2DEN, 2G3Q, 2MJ5, 2MRO, 2OOB, 2QHO, 2JY6, 3K9O, 4UN2, 6IF1, 6Q00, and 7F7X)^35–49^ were superimposed onto the TNK1-UBA domain structure to create a series of starting point TNK1-UBA:ubiquitin binding modes (PyMOL, Schrodinger). As the TNK1-UBA structure comprises two UBA-like motifs, each UBA:Ubiquitin crystal or NMR structure was superimposed onto both motifs, generating 33 unique candidate binding modes. The UBA and ubiquitin chains from these crystal and NMR structures were then replaced with the TNK1-UBA structure and a ubiquitin model from PDBID: 2MRO^41^. We also used AlphaFold2-Multimer^50^ to predict additional binding modes, using the TNK1-UBA structure and the ubiquitin structure from PDB ID: 2MRO^41^ as inputs. This generated an additional 250 candidate binding modes, each of which was then energy minimized using PyRosetta^51^. The difference in the solvent accessible surface area between the bound and unbound state of each model (ΔSASA)^52^, as well as the shape complementarity of each interface (SC)^53^, was then calculated. We also calculated the ΔSASA and SC of the UBA:ubiquitin interfaces of the fifteen wild type wwPDB structures used to generate some of our candidate binding modes (**Figure S2D**).

We clustered the 283 candidate binding modes into eight clusters based on the site of binding to the TNK1-UBA domain and compared each cluster to data from in vitro pull-down assays which used polyubiquitin and knockout mutants of the UBA domain. In brief, we incubated recombinant GST-UBA with K48- or K63-linked tetra-ubiquitin, captured the GST-UBA molecules using glutathione resin, and immunoblotted the captured proteins for both GST and ubiquitin. By comparing the pull-down data with our binding mode predictions, we observed that three out of eight potential binding mode clusters (**Figure S3**) were consistent with our pull-down data^6^ (**Figure 6A–B, Figure S3A–C)**. We rejected clusters whose binding mode either centered on amino acids whose substitution with aspartate did not abrogate ubiquitin binding, that did not involve the canonical binding amino acids of ubiquitin (L8, I44, H68, and V70), or that produced unresolvable clashes between the ubiquitin and the UBA domain. Furthermore, our previous data suggested that the F634 site is essential for binding to K48-but not K63-linked ubiquitin. An F634D substitution abrogated binding of K48-but not K63-linked ubiquitin^6^ (**Figure S3C**). This is not surprising because K48-linked ubiquitin adopts a compact conformation while K63-linked ubiquitin instead adopts an extended conformation^54^.

**Figure 6:**
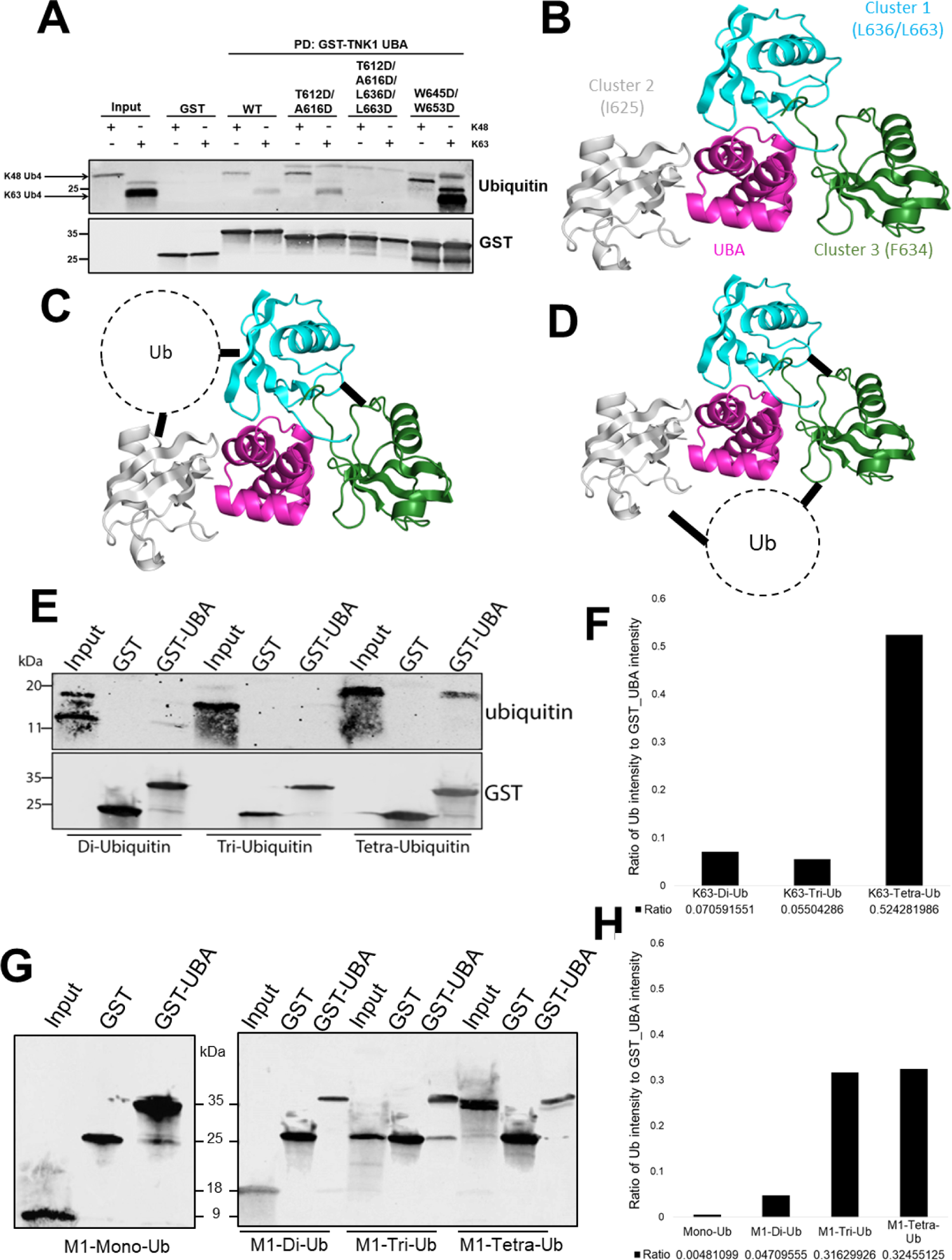
Predicted binding modes and corresponding experimental data. **A.** Western blot of TNK1-UBA mutants incubated with or without K48- or K63-tetra-ubiquitin. **B.** Predicted ubiquitin binding sites supported by in vitro pull-down data. **C-D.** Schematic of potential UBA:tetra-ubiquitin complexes. **E.** Western blot of TNK1-UBA incubated with K63-linked-di-, tri-, or tetra-ubiquitin. **F.** Quantified ratios between the GST-UBA and bound K63-ubiquitin. **G.** As in E, but with mono- or M1-linked-di-, tri-, or tetra-ubiquitin. **H.** As in in F, but with M1-linked-ubiquitins.

We were struck by the lack of consensus in precise ubiquitin orientation within many of the binding mode clusters, suggesting that some of the binding surfaces may bind ubiquitin only weakly, necessitating a highly avid interaction to maintain stable binding. Taken together, the modeling and experimental results suggest that the TNK1-UBA has multiple ubiquitin binding surfaces and would therefore be expected to bind polyubiquitin more tightly than mono-ubiquitin. One or more of the three proposed ubiquitin binding sites may be a higher-affinity anchor site that may guide polyubiquitin chains into stable binding interactions with the UBA domain. The additional bound ubiquitin molecules could then stabilize the first ubiquitin molecule and strengthen the affinity between the UBA and tetra-ubiquitin, explaining the remarkably low Kd’s previously observed for the TNK1-UBA interaction with K63- and K48-tetra-ubiquitin, (0.5 and 2.35 nM, respectively)^6^.

The insufficient length of mono-, di- and tri-ubiquitin may be the cause of their limited affinity because they fail to occupy the three UBA binding sites simultaneously. Since M1-tetra-ubiquitin gave a monodisperse SEC peak when bound to the UBA, we hypothesize that there may be at least one additional binding site for a fourth ubiquitin molecule, potentially either adjacent to between clusters 1 (cyan) and 2 (white) (**Figure 6C**) or between clusters 2 (white) and 3 (forest) (**Figure 6D**). In the first model, the ubiquitin would not be reasonably expected to interact with the UBA domain, consistent with our identification of three experimentally-validated binding sites.

Experiments confirm the UBA domain is selective for polyubiquitin chains. We repeated our pull-down experiments with mono, and K63- and M1-linked-di, tri, and tetra-ubiquitin. The UBA domain demonstrated significant binding to K63-tetra-ubiquitin, minimal binding to di- and tri-ubiquitin, and very minimal binding to mono-ubiquitin. Conversely, the UBA domain showed significant binding to M1-linked-tri- and tetra-ubiquitin, and minimal binding M1-linked-di-ubiquitin (**Figure 6E–H, S4**). These findings suggest that K63-linked-ubiquitin may tightly bind one or two sites on the UBA domain and weakly bind the remaining sites, requiring the avidity of a longer ubiquitin chain for stable binding. M1-linked-ubiquitin appeared to bind tightly to all three sites, allowing tri- and tetra-ubiquitin to function equally well. We hypothesize that the various linkages of polyubiquitin may have distinct binding modes to the UBA domain. For example, both the K63- and M1-ubiquitin molecules may occupy the same three sites on the UBA, but the K63-linkage may force one or more of the ubiquitin domains to bind using a less optimal binding orientation. These results provide evidence that the avidity of covalently linked ubiquitin chains and the specific linkage of ubiquitin molecules are crucial factors in achieving tight binding to the TNK1-UBA domain. Furthermore, the UBA domain can effectively assess the number of ubiquitin molecules and the types of ubiquitin linkages bound to it.

## Discussion

In this study, we provide a third example wherein 1TEL fusion enabled a protein to form crystals which diffracted to a resolution comparable to or better than the same protein crystallized on its own. In the current case, the TNK1-UBA domain formed crystals on its own as quickly as the 1TEL fusion, albeit with slightly less crystallization propensity, poorer crystal morphology (thin plates rather than hexagonal prisms), and generally poorer crystal quality (fraction of reflections indexed and mosaicity) (**Table S2**).

At the time that the first 1TEL-GG-UBA structure was solved, an accurate homology model was not available. Fortunately, the 1TEL domain constituted a sufficient fraction of the atoms in the asymmetric unit to solve the crystallographic phases adequate to build in the UBA domain. The ability to use TELSAM as a molecular replacement search model may however see limited usefulness, as it is currently limited to proteins smaller than the TELSAM variant used and sufficiently accurate predicted structures are now available for the majority or proteins^31^.

The unprecedented discovery that 1TEL–UBA formed crystals at protein concentrations as low as 0.1 mg/mL promises to enable the crystallization and high-resolution structure determination of proteins exhibiting limited solubility or that can only be practically produced in ug quantities. This also suggests that TELSAM-fusions may form increasingly smaller protein crystals as the protein concentration is lowered. The ability to predictably form very small crystals at very low protein concentrations offers a potential solution to a significant problem in microelectron diffraction (microED) studies of protein and small molecule crystals: Because electrons interact with matter far more readily than do X-rays, an electron passing through a crystal has a high likelihood of interacting with more than one atom before exiting the crystal (dynamical scattering). This likelihood increases the larger the crystal, setting an upper limit of crystal thickness for microED of 100–200 nm^55^. Crystals in this size range cannot be detected in crystal drop using visible or UV light microscopy or second harmonic generation detection, requiring researchers to transfer drops containing larger visible crystals to cryo-electron microscopy grids and then search those grids for microcrystals of usable size. Current solutions to this problem include gentle sonication of mosaic protein crystals into smaller fragments^22^ or focused ion beam (FIB) milling of larger crystals down to the desirable size^56^. These techniques are either unreliable (sonication) or require additional equipment and expertise and are time intensive (FIB milling), resulting in a significant barrier to the average crystallographer or electron microscopist. TELSAM’s ability to predictably form crystals in the 100–200 nm size range could shatter these barriers and is currently under investigation.

This study provides structural insight into the selectivity of the TNK1-UBA domain for polyubiquitin, and also raises questions about the role of this UBA domain in TNK1 function and regulation. Interestingly, TNK1 and its sister kinase ACK1 are, to our knowledge, the only examples of kinases with functional UBA domains^6,57, 58^. Our previous data suggest that loss of the UBA domain alters the substrate profile of TNK1^6^, so it is possible that interactions between the TNK1 UBA and polyubiquitin tether the kinase to substrates. It is also perhaps likely that the interaction between the TNK1 UBA and ubiquitinated species results in TNK1 protein turnover, given that a naturally occurring C-terminal truncation of TNK1 in Hodgkin lymphoma, which deletes the UBA domain, results in high levels of TNK1 protein^2^. Another intriguing question relates to how inhibitory 14-3-3 binding regulates the UBA domain of TNK1. The 14-3-3 binding site in TNK1 sits immediately adjacent to the UBA domain, suggesting that 14-3-3 may inhibit TNK1 by masking the UBA domain. Additional work, including co-structures of TNK1 with 14-3-3, will help answer these questions.

## Acknowledgements

This study was supported by an NIH R15 GM146209 to JDM and an NIH R01 GM147310-01 to JLA. CME was supported by graduate fellowships from the Simmons Center for Cancer Research. Use of the Stanford Synchrotron Radiation Lightsource, SLAC National Accelerator Laboratory, is supported by the U.S. Department of Energy, Office of Science, Office of Basic Energy Sciences under Contract No. DE-AC02-76SF00515. The SSRL Structural Molecular Biology Program is supported by the DOE Office of Biological and Environmental Research, and by the National Institutes of Health, National Institute of General Medical Sciences (P30GM133894). The contents of this publication are solely the responsibility of the authors and do not necessarily represent the official views of NIGMS or NIH.

## Author contributions

Conceptualization, J.A. and J.M.; Methodology, S.N., S.S., Y.T., J.A., and J.M.; Formal Analysis, Y.T. and J.M.; Investigation, S.N; Y.T., S.S., T.S., M.P., W.A., C.E., D.M., D.B., B.W, E.T., C.S., and S.B.; Resources, T.D., J.A., J.M.; Writing – Original Draft, S.N., Y.T., S.S., T.S., M.P., J.A, and J.M.; Writing – Review & Editing, Y.T., T.D., J.A., and J.M.; Visualization, S.N., Y.T., T.S., C.E., D.M., and J.M; Funding Acquisition, J.A. and J.M.; Supervision, J.A. and J.M.

## STAR★Methods

### Key resources table

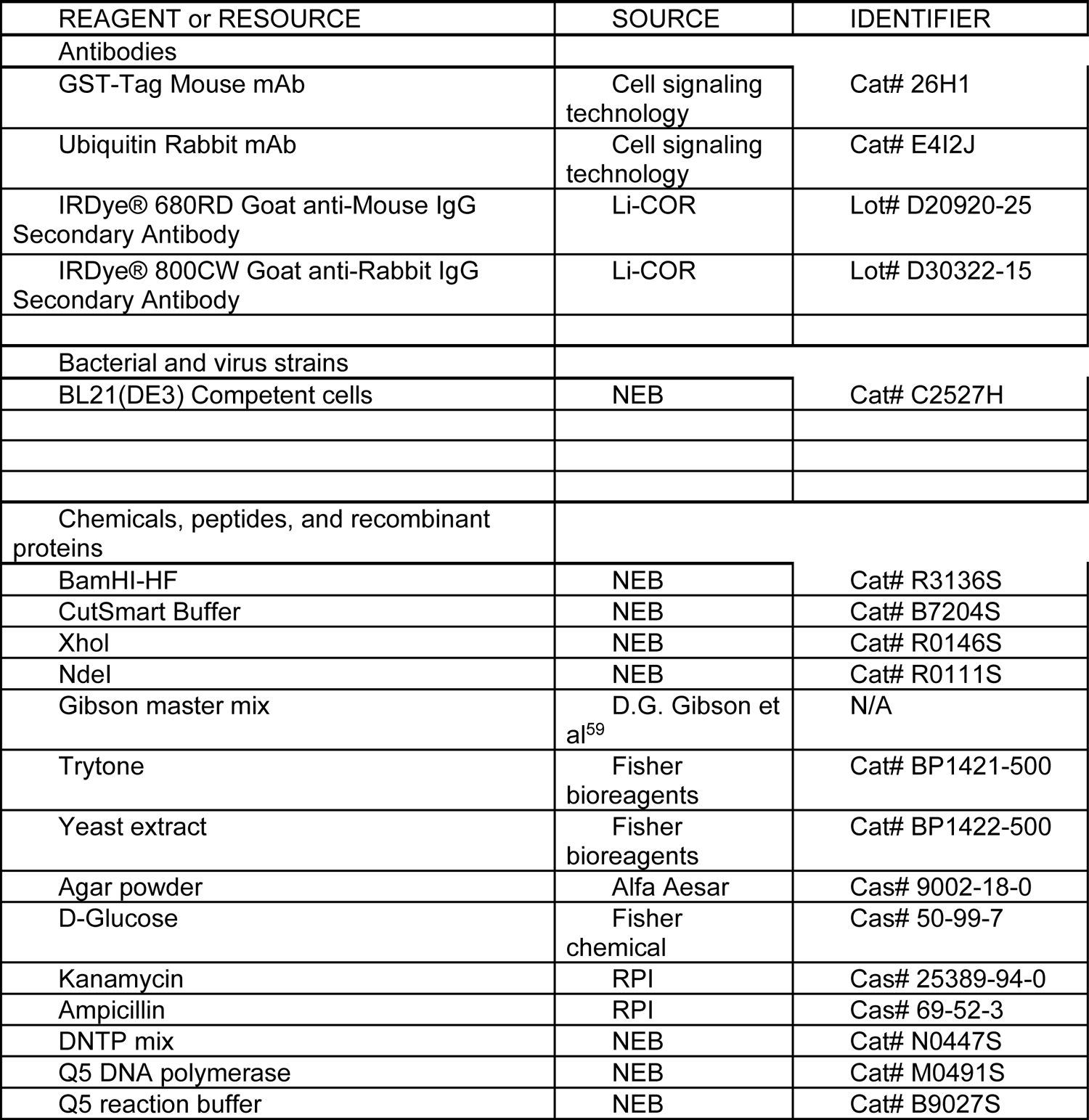
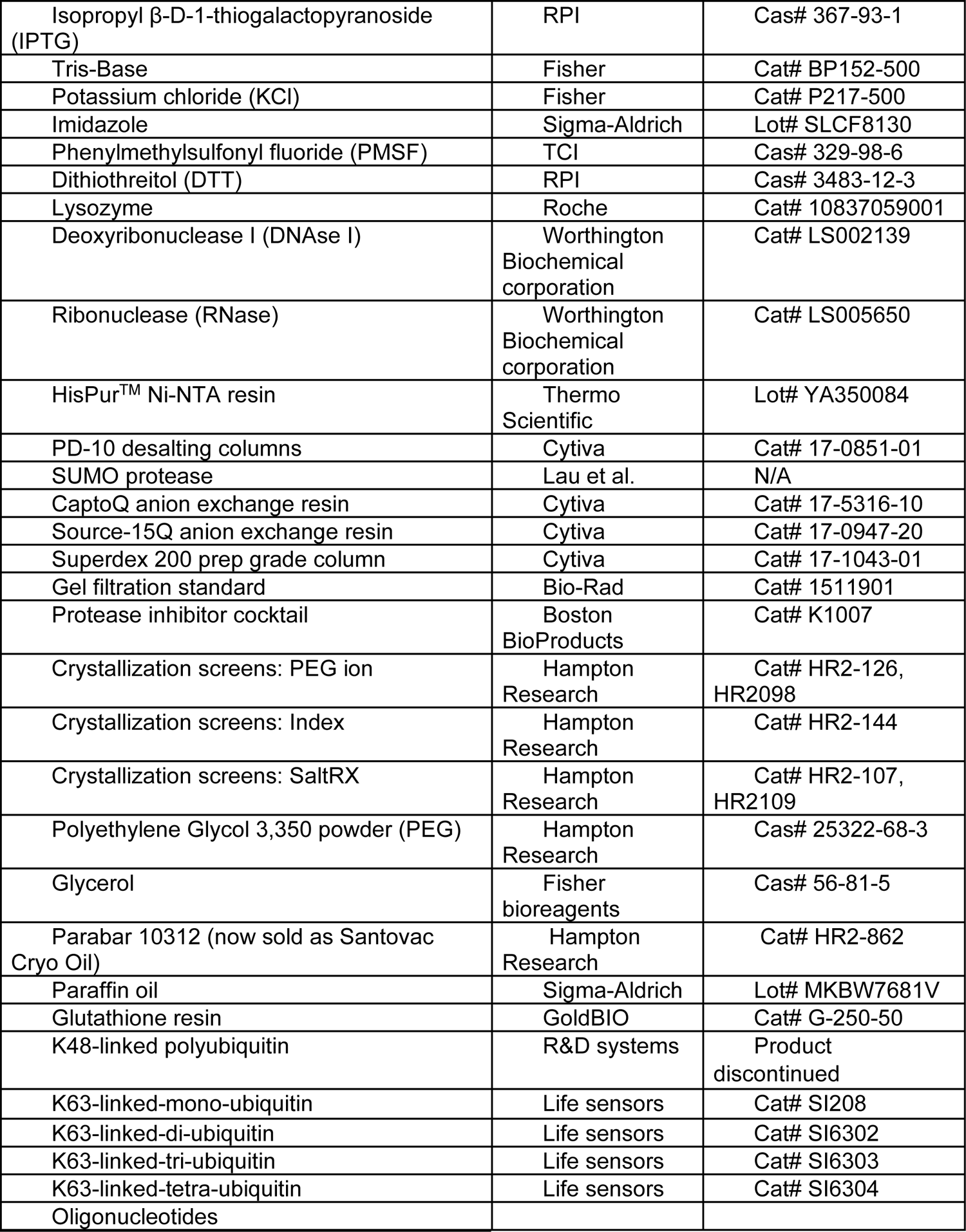
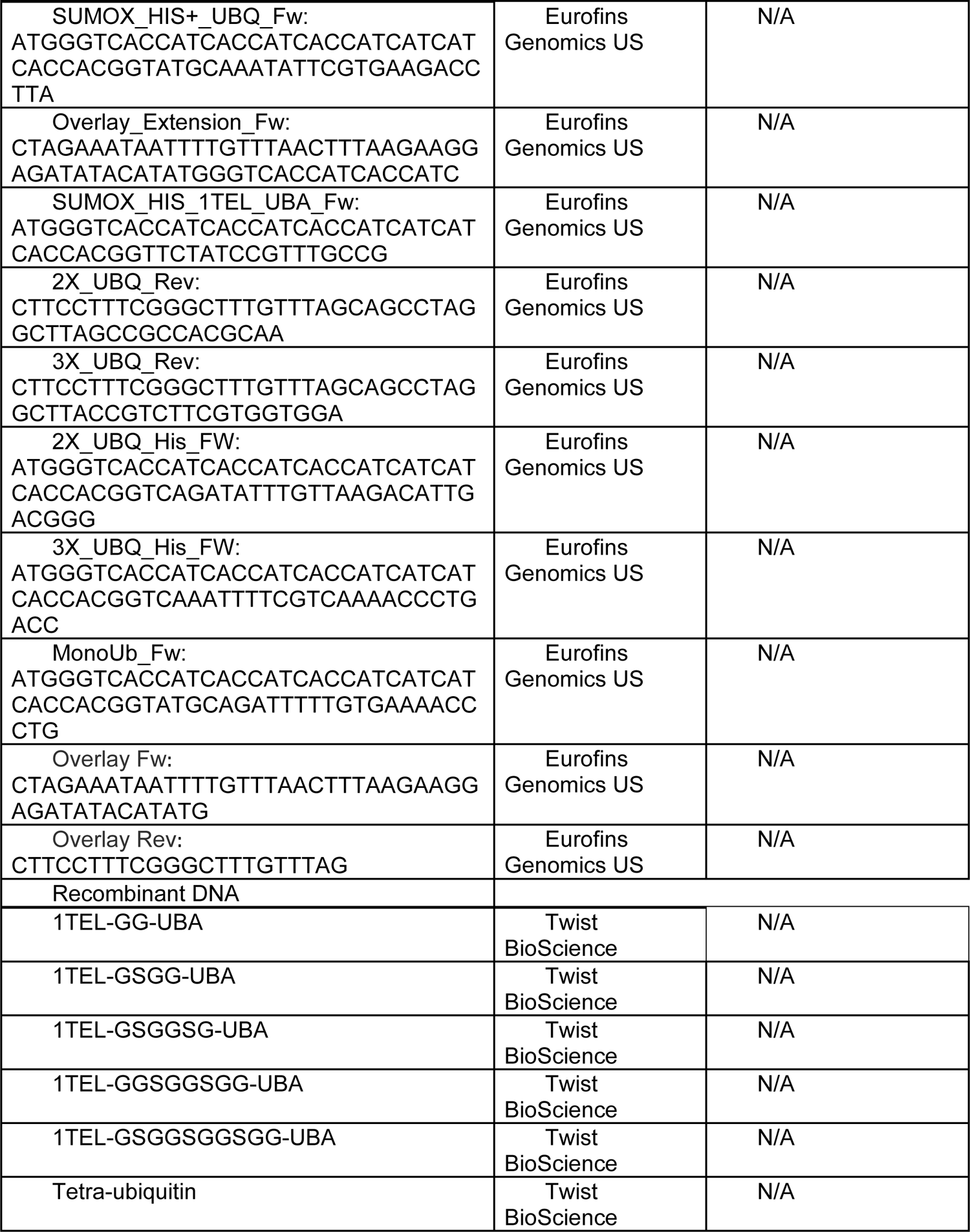
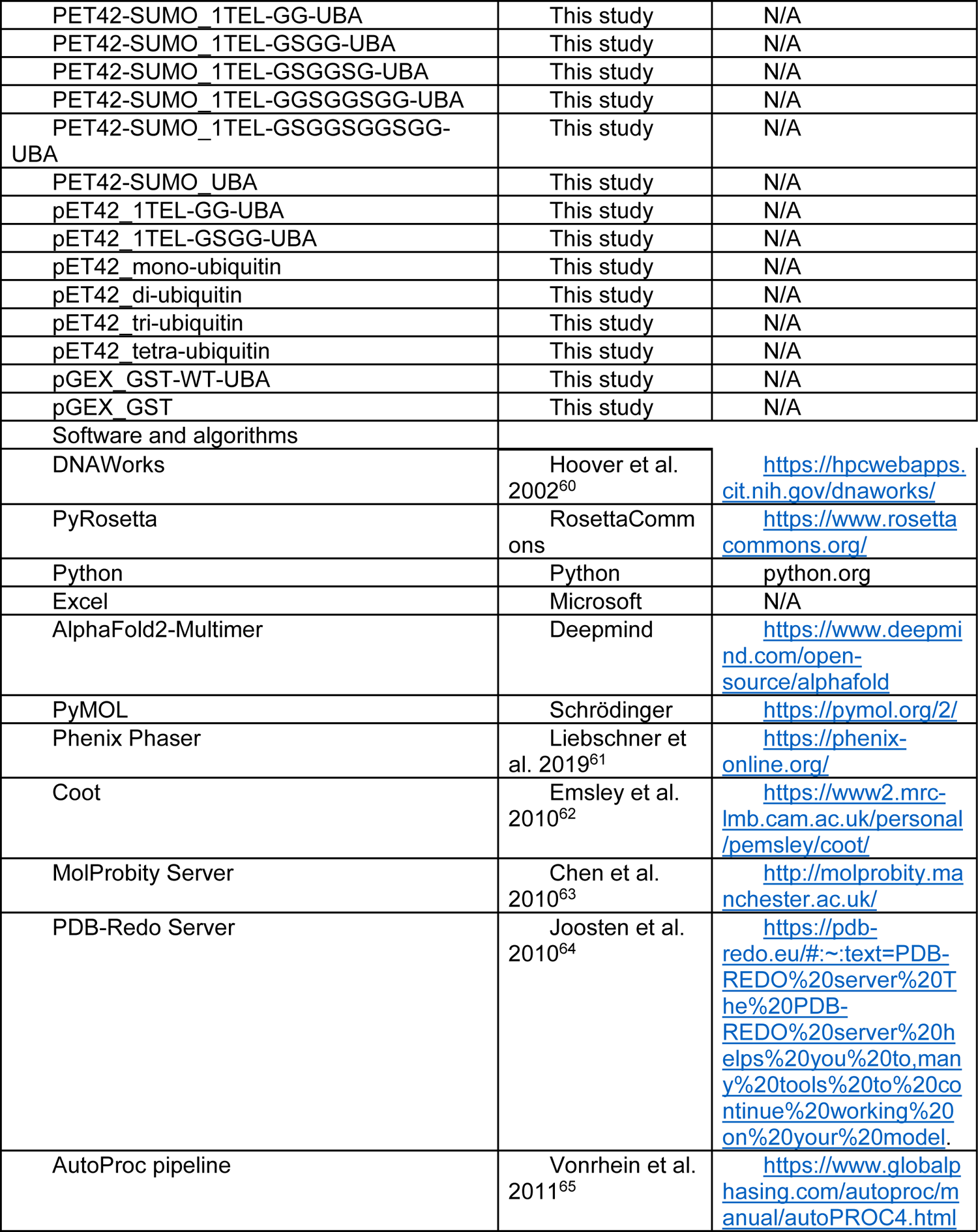
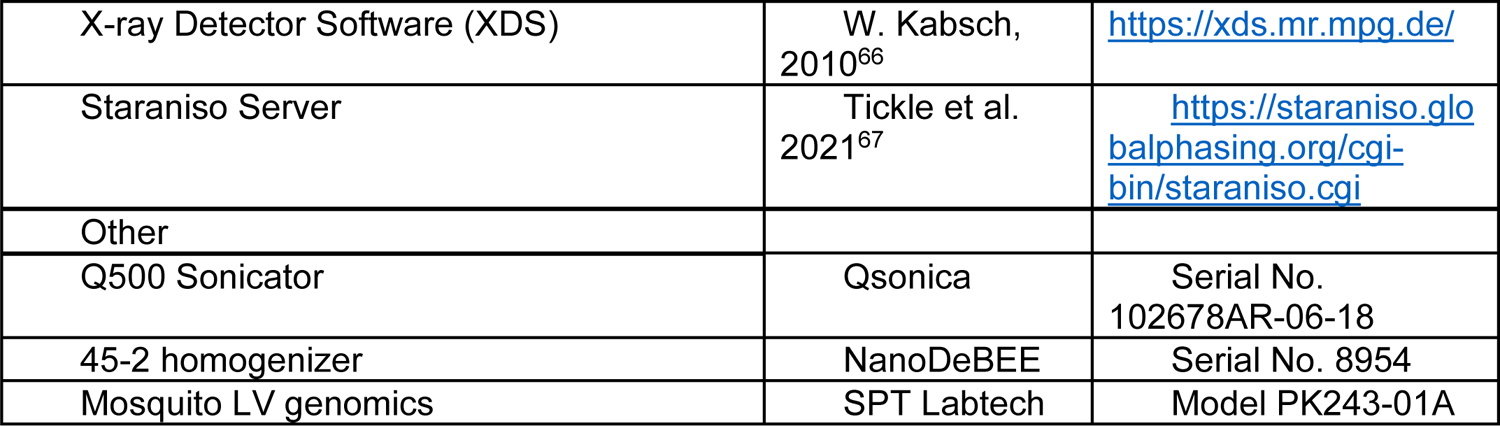

#### Resource availability Lead contact

Further information and requests for resources and reagents should be directed to and will be fulfilled by the lead contact, James D. Moody.

### Materials availability

Most of the materials used in this study are publicly or commercially available. All unique and stable reagents generated in this study are available from the lead contact without restriction, upon reasonable request.

### Data and code availability

All data are available from the lead contact upon reasonable request.

All algorithms used in this study are publicly available. The purpose-written software has been deposited at https://github.com/yijietseng/General_scripts/blob/main/relax.py and https://github.com/yijietseng/General_scripts/blob/main/measure_sc_buried_sa.py and is publicly available.

Any additional information required to reanalyze the data reported in this work paper is available from the lead contact upon request.

### Method details

#### Cloning of the TNK1-UBA domain alone

Residues 590–666 of human Thirty-eight Negative Kinase-1 (TNK1, Uniprot Q13470) were reverse-translated, codon-optimized (DNAworks)^60^, and inserted downstream of the 10xHis-SUMO domain in the pET42-SUMO vector^68^. Both cysteines in the UBA domain were substituted with alanines to prevent oxidation-mediated misfolding and aggregation due to unwanted disulfide bond formation. A gene fragment containing the UBA domain was synthesized by a commercial vendor (Twist Biosciences), assembled into the pET42-SUMO vector^68^ using Gibson assembly^59^, introduced into BL21(DE3) cells, and sequence verified (Eton Biosciences).

#### Cloning of 1TEL-GG-UBA

Residues 590–666 of human Thirty-eight Negative Kinase-1 (TNK1, Uniprot Q13470) were inserted after residues 48–123 of human ETV6/TEL (the sterile alpha motif/SAM domain, Uniprot P41212), both of which were inserted downstream of the 10xHis-SUMO domain in the pET42-SUMO vector^68^. Residues 122–123 of the TELSAM domain were substituted with glycines to impart flexibility to the connection. Other substitutions in the TELSAM domain included V112E, critical for pH-triggerable polymerization, as well as R80S (carried over from the structure of the TELSAM domain used in the modeling (PDBID: 2QAR)^30^. Both cysteines in the UBA domain (C610 and C644) were substituted with alanines. This construct was synthesized, cloned, and sequence verified as described above.

#### Cloning of 1TEL-GSGG-UBA, 1TEL-GSGGSG-UBA, 1TEL-GGSGGSGG-UBA, and 1TEL-GSGGSGGSGG-UBA

To make the 1TEL-GSGG-UBA, 1TEL-GSGGSG-UBA, 1TEL-GGSGGSGG-UBA, and 1TEL-GSGGSGGSGG-UBA constructs, the number of residues between the TEL-SAM domain and the UBA were increased to 4, 6, 8, and 10, respectively, using a repeating “SGG” pattern. These constructs were synthesized, cloned, and sequence verified as described above.

#### Cloning of SUMO tag-free 1TEL-GG-UBA and 1TEL-GSGG-UBA

To avoid using SUMO protease in later experiments, a 1TEL-GG-UBA construct without the SUMO domain was synthesized by PCR amplifying the 1TEL-GG-UBA domain from the existing 10xHis-1TEL-GG-UBA construct. PCR was also used to attach a 10xHis tag directly to the N-terminal end of the TELSAM domain. To prepare a pET42-derived vector without the SUMO domain, the 10xHis-SUMO domain was removed by restriction digest at the XhoI and NdeI cut sites. The 10xHis-1TEL-GG-UBA fragment was assembled into the prepared pET42-derived vector and sequence verified as described above.

#### Cloning of Mono- and M1-linked Di-, Tri- and Tetra-Ubiquitin

The initial tetra-ubiquitin construct was created by extracting the M1-di-ubiquitin amino acid sequence of human polyubiquitin (Uniprot P0CG47) and reverse translating it to make the DNA sequence of each of the ubiquitin repeats as unique as possible. Two gene fragments containing the di-ubiquitin were synthesized by Twist Biosciences, the first having a 3’ XhoI site. The first M1-di-ubiquitin fragment was then assembled into the pET42-SUMO vector and sequence verified as described above. The sequence-verified vector was then digested with XhoI and the second M1-di-ubiquitin fragment assembled into it, followed by sequence verification. We later used this construct as a template to create SUMO tag-free M1-linked mono-, di-tri- and tetra-ubiquitin fragments via PCR, appending a 10xHis tag in each case. Each fragment was assembled into the NdeI/XhoI digested pET42-derived vector and sequence verified as described above.

#### Protein expression and purification

First, 40-60 mL of Luria-Bertani (LB) media, supplemented with 0.35% glucose and 100 µg/mL kanamycin, were inoculated with a stab of frozen cell stock and shaken at 30°C and 250 RPM for 16 hours. The next day, 10 mL of this overnight culture was diluted into one liter of LB media supplemented with 0.05% glucose and 100 μg/mL kanamycin. The media was shaken again at 37°C and 250 rpm until it reached an O.D. of 0.5. At this O.D., IPTG was added to a final concentration of 100 µM. The culture was shaken at 18°C and 250 RPM for 20-24 more hours, and the cells were collected using centrifugation, snap-frozen in liquid nitrogen, and stored at −80°C.

All protein purification steps were performed in a 4°C refrigerator or on ice. First, 5-20 g of wet cell paste was resuspended in 5 cell paste volumes of wash buffer (50 mM Tris, pH 8.8, 200 mM KCl, 50 mM imidazole), supplemented with 1 mM phenylmethylsulfonyl fluoride (PMSF) and 100 μM dithiothreitol (DTT). Then, final concentrations of 20 μM Lysozyme, 800 nM deoxyribonuclease I, and 2 μM ribonuclease were added to the resuspended cells. The cells were lysed using sonication for 25 cycles of 12 seconds on, 59 seconds off at 60% power (Qsonica Q500). Cell suspensions of SUMO tag-free 1TEL-GG-UBA and ubiquitin variants were instead lysed at 18,000 psi in 2 passes through a NanoDeBEE 45-2 homogenizer. In each case, the resulting lysate was clarified by centrifugation at 40,000 x g and applied to 2–3 mL of HisPure Ni-NTA resin (Thermo). The resin was then washed with 7-20 column volumes (CV) of wash buffer. After that, the protein was eluted with around 7-10 CV of elution buffer (50 mM Tris, pH 8.8, 200 mM KCl, 400 mM imidazole) and desalted using several PD-10 desalting columns in parallel (Cytiva). For one liter of cell culture, typical protein yields were 120 mg for the UBA-alone, 80 mg for 1TEL-GG-UBA, 100 mg for 1TEL-GG-UBA, 1TEL-GSGGSG-UBA, 1TEL-GGSGGSGG-UBA, and 1TEL-GSGGSGGSGG-UBA, 189 mg for M1-tetra-ubiquitin, 152 mg for M1-tri-ubiquitin, 120 mg for M1-di-ubiquitin, 150 mg for M1-mono-ubiquitin, and 50 mg of SUMO-free 1TEL-GG-UBA and 1TEL-GSGG-UBA. The SUMO tag was removed from those constructs that included it by adding 0.1-0.5 mg SUMO protease^69^ and DTT to 0.1 mM. The cleavage reaction proceeded overnight at 4°C. The protein solution was then passed over 2 mL of fresh Ni-NTA resin to remove the cleaved SUMO tags and SUMO protease.

After being diluted 8-fold with water, the 1TEL-GSGG-UBA and 1TEL-GSGGSG-UBA were applied to 4 mL of either CaptoQ or Source-15Q anion exchange resin (Cytiva). These proteins bound to the anion exchange resin and were eluted in a KCl gradient. Size exclusion chromatography was used to further purify all the proteins using a 100 mL Superdex 200 prep grade column (Cytiva), which was calibrated using a gel filtration standard (Bio-Rad). After SEC, the proteins were buffer exchanged into 12.5 mM Tris, pH 8.8, 200 mM KCl. A cocktail of 6 protease inhibitors (100 mM AEBSF, 80µM Aprotinin, 1.5 mM E-64, 2 mM Leupeptin, 1 mM Pepstatin A, 5 mM Bestatin) (Boston BioProducts) was added to each TELSAM fusion before setting crystal trays.

#### Crystallization and diffraction of 1TEL-GG-UBA

1.2 μL of 0.1, 0.2, 0.5, 1, 2, 5, 10, and 15 mg/mL 1TEL-GG-UBA was combined with 1.2 μL of reservoir solution in a sitting drop format (SPT Labtech Mosquito). Commercially available crystallization screens (PEG Ion, Index, and Salt-RX) (Hampton Research) and custom screens (PEG-custom) were used. Crystals appeared in three days in 3 distinct crystallization conditions. From these, a set of 96 BisTris Mg-Formate optimization conditions were created and screened. The largest crystals (250 μm) appeared in 100 mM Bis-Tris pH 7.0, 200 mM Mg-formate. Before freezing in liquid nitrogen, the crystals were mounted using either 20% glycerol in crystallization reservoir solution as a cryo-protectant or a 50:50 Parabar 10312:Paraffin oil mixture. No further optimization of cryoprotectants or freezing conditions was carried out. X-ray diffraction data was collected remotely at SSRL beamline 9-2. These crystals diffracted to 1.53–2.70 Å resolution (19 crystals, I/σ ≥ 2) and were readily indexed in a primitive hexagonal unit cell with dimensions a = b = 67.1–68.2 Å, c = 55.5–61.4 Å. The crystals exhibited low estimated mosaicity (0.13–1.50°) and indexed 16–89% of non-ice reflections.

#### Crystallization and diffraction of 1TEL-GSGG-UBA, 1TEL-GSGGSG-UBA, 1TEL-GGSGGSGG-UBA, and 1TEL-GSGGSGGSGG-UBA

These constructs were crystallized at a concentration of 20 mg/mL as described above. Commercially available crystallization screens (PEG Ion, Index, and Salt-RX [Hampton Research]) and custom screens (PEG-custom and BisTris Mg-Formate) were used. Crystals of 1TEL-GSGG-UBA appeared in 10 conditions in 10–14 days. The largest (150 μm) appeared in 100 mM Bis-Tris-propane, pH 7.0, 2.5 M Ammonium nitrate. Crystals were also seen in conditions containing Ammonium Chloride and in Bis-Tris Mg-formate. The crystals were mounted, frozen, and diffraction data was collected as described above. These crystals diffracted to 1.89–2.27 Å resolution (4 crystals, I/σ ≥ 2) and were readily indexed in a primitive hexagonal unit cell with dimensions a = b = 67.2–68.8 Å, c = 54.7–59.6 Å. The crystals exhibited moderate estimated mosaicity (0.36–1.50°) and indexed 59–79% of non-ice reflections. Highly non-merohedrally twinned crystals of 1TEL-GSGGSG-UBA were observed after 85 days in 0.2 M Calcium chloride dihydrate, pH 5.1, 20% w/v Polyethylene glycol 3,350. These crystals did not diffract X-rays. The 1TEL-GGSGGSGG-UBA and 1TEL-GSGGSGGSGG-UBA constructs did not yield crystals, even after several months.

#### Crystallization of the TNK1-UBA-alone

The UBA-alone was crystallized at a concentration of 3.1 mg/mL as described above. Commercially available crystallization screens (PEG Ion, Index (Hampton Research), and custom screens (PEG-custom and BIS-TRIS magnesium formate) were used. Two days after setting the trays, square shaped plate crystals (250 μm) appeared in 2 conditions (20–40 mM citric acid, 60–80 mM Bis-Tris-propane, pH 6.4–8.8, 16–20% PEG 3350). The crystals were mounted, frozen, and diffraction data was collected as described above. These crystals diffracted to 1.38–1.74 Å resolution (3 crystals, I/σ ≥ 2) and were readily indexed in a primitive orthorhombic unit cell with dimensions a = 53.3–53.7 Å, b = 87.8–88.1 Å, c = 26.2–26.5 Å. The crystals exhibited moderate estimated mosaicity (0.3–1.3°) and indexed 20–64% of non-ice reflections.

#### Data reduction and structure solution

The Autoproc pipeline^65^ was used to process the datasets and the Staraniso algorithm was used to estimate the anisotropic diffraction limits^67^. Molecular replacement using Phenix Phaser^61, 70^ was employed to solve the phases. The structures then went through alternating stages of rebuilding in Coot^62^ and refinement in Phenix Refine^71, 72^. TLS groups calculated by Phenix were used to refine TLS parameters as well. Refinement was assisted using statistics from the MolProbity server^63^. The PDB-Redo server was used to assist refinement of the 1TEL-GSGG-UBA and the 1TEL-GG-UBA (2 mg/mL)^64^.

#### Computational binding mode prediction and score calculations

wwPDB-derived TNK1-UBA:Ubiquitin binding mode models were constructed as described in the main text. The amino acid sequences of TNK1-UBA and mono-, di-, tri-, and tetra-ubiquitin were also submitted to AlphaFold2-Multimer predictions using the research supercomputing resources at Brigham Young University. The resulting structures were optimized using the FastRelax algorithm via PyRosetta^51^. The difference in solvent accessible surface area between the bound and unbound state (ΔSASA)^52^ and the shape complementary (SC)^53^ between TNK1-UBA and each ubiquitin subunit were then calculated using Python and PyRosetta.

#### UBA:ubiquitin pull-down assay

GST-tagged TNK1-UBA domain was produced as previously described^6^. In brief, recombinant GST-TNK1-UBA protein (WT and mutants) were pulled down using glutathione resin (GoldBIO). The pulled down GST-tagged proteins were incubated individually with K48-(R&D systems) or K63-linked-tetra-ubiquitin (Life sensors) on a rotor for one hour at 4°C. Following that, the mixtures were subjected to a wash step using a wash buffer composed of 10 mM HEPES at pH 7.5, 300 mM NaCl, and 1 mM DTT. The resin, which contained the GST-tagged protein along with the captured ubiquitin, was subsequently boiled, run out by SDS-PAGE, and immunoblotted using antibodies against GST and ubiquitin (Cell Signaling Technology). Furthermore, to conduct experiments on the selectivity of the UBA domain towards polyubiquitin, we incubated 1 µg of K63- and 2 µg of mono or M1-linked-di-, tri-, and tetra-ubiquitin, followed by pulldown and immunoblot using the procedure described.

## Supplementary data

**Table S1.** Crystallographic Data Collection and Refinement Statistics

| Construct | UBA-alone | 1TEL-GG-UBA<br>(15 mg/mL) | 1TEL-GG-UBA<br>(2 mg/mL) | 1TEL-GG-UBA<br>(0.5 mg/mL) | 1TEL-GSGG-UBA<br>(20 mg/mL) |
| --- | --- | --- | --- | --- | --- |
| PDBID | 7TCY | 7U4W | 7TDY | 7U4Z | 7T8J |
| **Data Collection** |  |  |  |  |  |
| X-ray source | SSRL<br>BEAMLINE BL9-2 | SSRL<br>BEAMLINE BL9-2 | SSRL<br>BEAMLINE BL9-2 | SSRL<br>BEAMLINE<br>BL9-2 | SSRL BEAMLINE<br>BL9-2 |
| Wavelength | 0.979460 | 0.979460 | 0.979460 | 0.979460 | 0.979460 |
| Detector Type | DECTRIS<br>PILATUS 6M | DECTRIS<br>PILATUS 6M | DECTRIS<br>PILATUS 6M | DECTRIS<br>PILATUS 6M | DECTRIS<br>PILATUS 6M |
| Detector<br>Distance (mm) | 300 | 300 | 300 | 200 | 300 |
| Resolution range | 34.06–1.54<br>(1.60–1.54)* | 40.48–2.10<br>(2.18–2.10) | 29.43–1.53<br>(1.59–1.53) | 33.69–2.03<br>(2.10–2.03) | 34.33–1.89<br>(1.96–1.89) |
| Space group | P 21 21 2 | P 65 | P 65 | P 65 | P 65 |
| Cell a, b, c (Å) | 53.734, 88.070,<br>26.497 | 67.971, 67.971,<br>55.749 | 67.221, 67.221,<br>60.888 | 67.375 67.375<br>58.683 | 68.651, 68.651,<br>56.735 |
| Cell $\alpha$, $\beta$, $\gamma$ (°) | 90, 90, 90 | 90, 90, 120 | 90, 90, 120 | 90, 90, 120 | 90, 90, 120 |
| Total reflections | 189390 (15901) | 123586 (11687) | 423148 (35018) | 172181 (19841) | 170711 (17626) |
| Unique<br>reflections | 18888 (1846) | 8852 (820) | 22513 (2344) | 8835 (976) | 12236 (1208) |
| Multiplicity | 10.0 (8.6) | 14.5 (14.1) | 18.8 (14.9) | 19.5 (20.2) | 14.0 (14.6) |
| Completeness<br>(%) | 97.62 (99.51) | 98.32 (96.24) | 95.11 (99.96) | 89.42 (100.00) | 99.92 (100.00) |
| I/σI | 19.86 (2.48) | 28.14 (2.27) | 27.17 (2.69) | 11.07 (2.39) | 20.93 (2.59) |
| Wilson B-factor | 21.01 | 51.29 | 29.30 | 27.18 | 39.92 |
| R-merge | 0.0557 (0.813) | 0.0408 (1.08) | 0.0488 (1.03) | 0.220 (1.84) | 0.06456 (1.63) |
| R-meas | 0.0588 (0.865) | 0.0423 (1.13) | 0.0502 (1.07) | 0.226 (1.89) | 0.0670 (1.69) |
| R-pim | 0.0184 (0.289) | 0.0111 (0.297) | 0.0114 (0.273) | 0.0511 (0.419) | 0.0179 (0.442) |
| CC1/2 | 0.999 (0.882) | 1.000 (0.923) | 0.999 (0.907) | 0.998 (0.839) | 0.999 (0.903) |
| CC* | 1.000 (0.968) | 1.000 (0.98) | 1.000 (0.975) | 0.999 (0.955) | 1.000 (0.974) |
| **Refinement** |  |  |  |  |  |
| Reflections used in refinement | 18888 (1846) | 8493 (818) | 22504 (2343) | 8827 (980) | 12228 (1208) |
| Reflections used for R-free | 993 (85) | 467 (33) | 1080 (119) | 430 (42) | 595 (49) |
| R-work | 0.1886 (0.2954) | 0.2452 (0.3866) | 0.2052 (0.2893) | 0.2016 (0.2760) | 0.2238 (0.3423) |
| R-free | 0.2134 (0.3711) | 0.2756 (0.3834) | 0.2176 (0.3138) | 0.2318 (0.3466) | 0.2348 (0.3845) |
| CC (work) | 0.962 (0.891) | 0.956 (0.749) | 0.959 (0.866) | 0.957 (0.846) | 0.955 (0.844) |
| CC (free) | 0.970 (0.753) | 0.885 (0.689) | 0.942 (0.824) | 0.965 (0.849) | 0.948 (0.736) |
| Number of non-H atoms | 1384 | 1070 | 1238 | 1237 | 1115 |
| Macromolecule atoms | 1237 | 1061 | 1150 | 1196 | 1095 |
| Ligands | 24 | 0 | 24 | 1 | 1 |
| Water | 127 | 9 | 70 | 40 | 19 |
| Protein residues | 154 | 144 | 148 | 153 | 147 |
| Bond length R.M.S. Deviations | 0.011 | 0.004 | 0.010 | 0.005 | 0.003 |
| Bond angle R.M.S. Deviations | 1.20 | 0.57 | 1.12 | 0.62 | 0.54 |
| **Ramachandran Stats (%)** |  |  |  |  |  |
| Favored (%) | 99.33 | 98.57 | 100.00 | 98.68 | 97.20 |
| Allowed (%) | 0.67 | 1.43 | 0.00 | 1.32 | 2.80 |
| Outliers (%) | 0.00 | 0.00 | 0.00 | 0.00 | 0.00 |
| Rotamer outliers (%) | 0.00 | 0.00 | 0.00 | 0.00 | 0.00 |
| Clashscore | 2.01 | 2.04 | 4.44 | 0.00 | 2.88 |
| **B-factors (Å²)** |  |  |  |  |  |
| Average B-factor | 29.26 | 72.80 | 40.22 | 33.07 | 56.32 |
| Macromolecules | 28.21 | 72.86 | 39.78 | 33.02 | 56.42 |
| Ligands | 47.38 | NA | 52.00 | 65.02 | 64.02 |
| Water | 36.60 | 65.30 | 44.42 | 33.70 | 50.59 |
| Number of TLS groups | 11 | 6 | 9 | 7 | 11 |

\*Statistics for the highest-resolution shell are shown in parentheses.

**Table S2.**
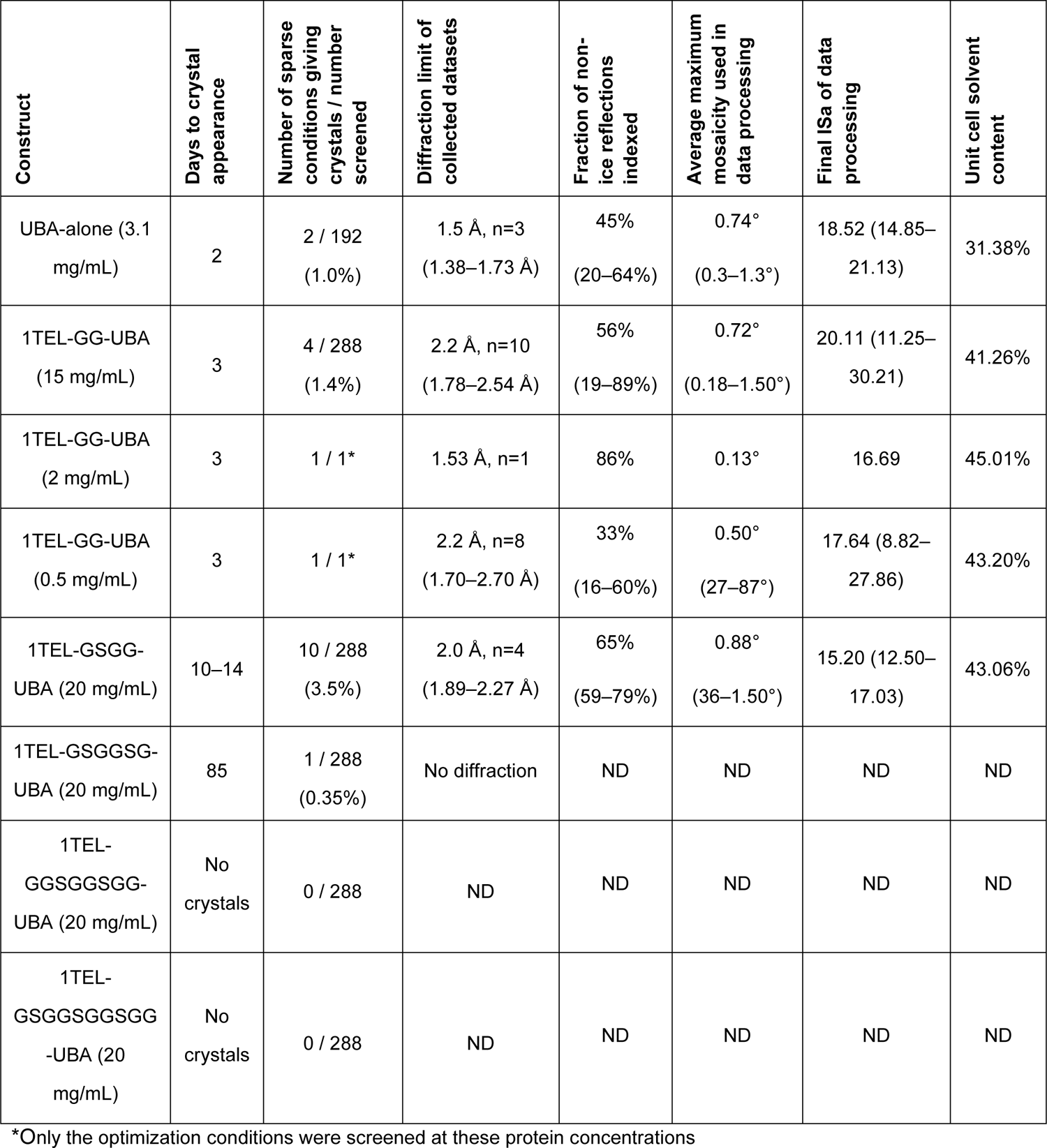
Crystallization Time, Propensity, and Diffraction Quality of UBA Constructs

**Figure S1:**
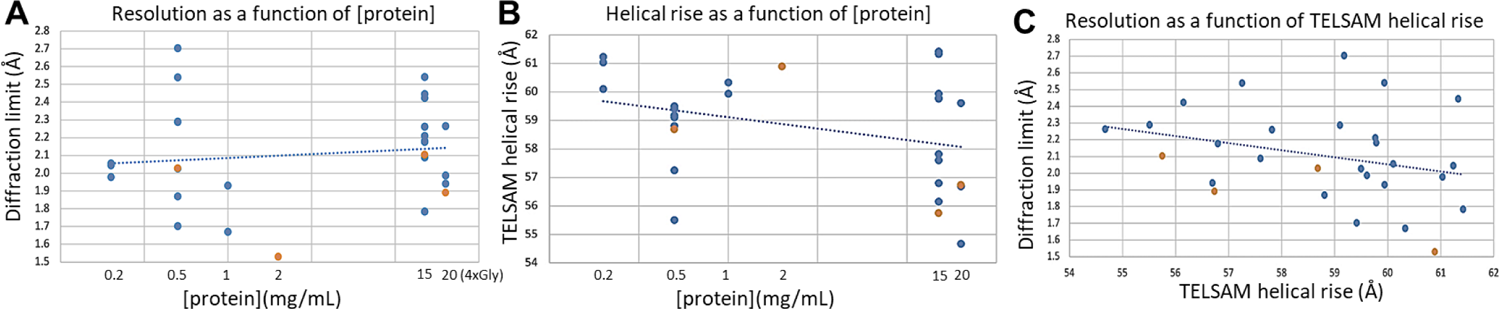
Comparison between diffraction resolution, protein concentration and helical rise. **A.** Resolution as a function of protein concentration. **B.** Helical rise as a function of protein concentration. **C.** Resolution as a function of TELSAM helical rise.

**Figure S2:**
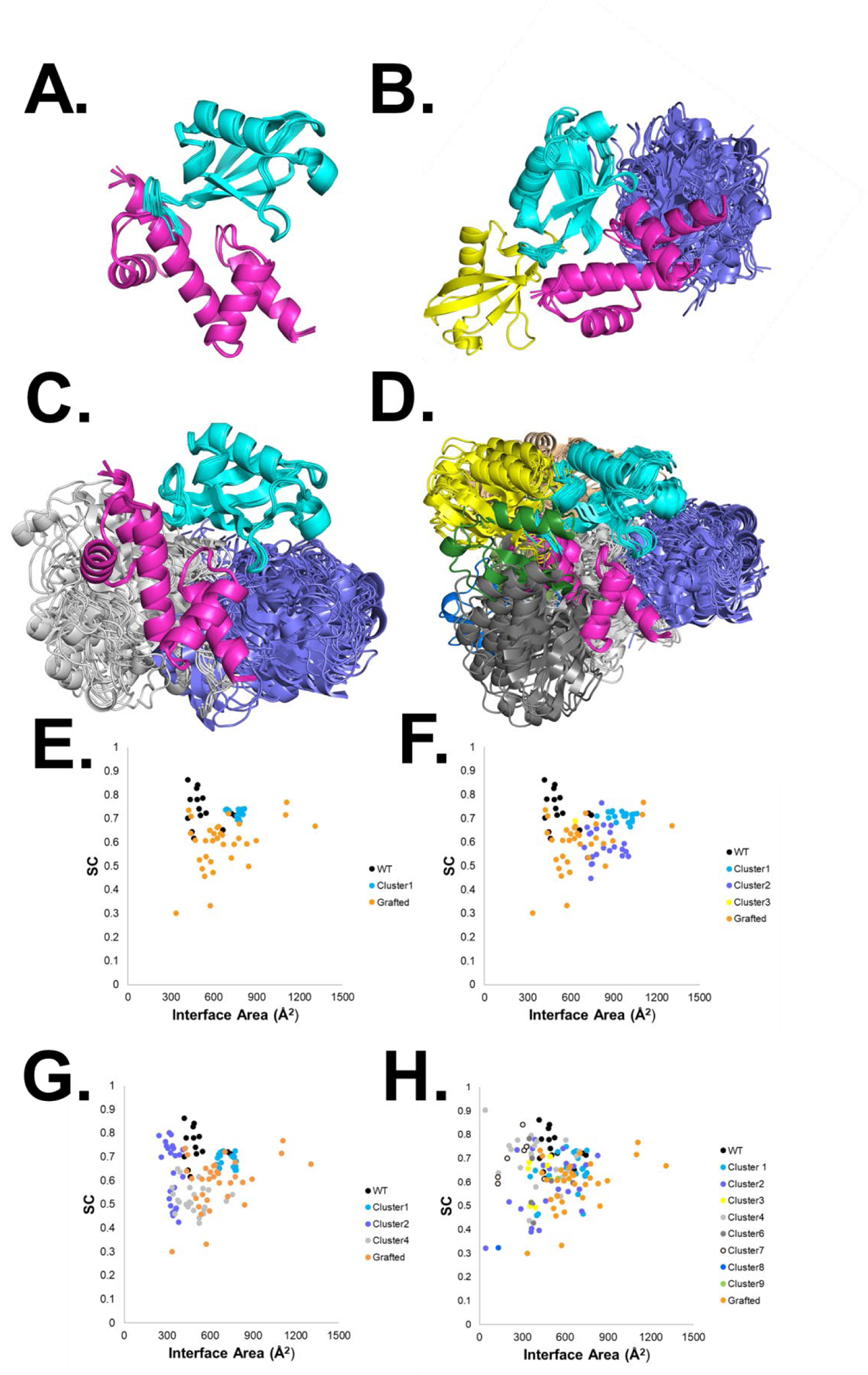
UBA:Ubiquitin binding site Clusters identified by AlphaFold2-Multimer: **A-D.** AlphaFold2-Multimer predicted models using mono-, di-, tri-, and tetra-ubiquitin (cyan, purple, yellow, forest, white, dark gray) and the TNK1 UBA (magenta). **E-H.** Distribution of the interface areas and SC of individual models.

**Figure S3:**
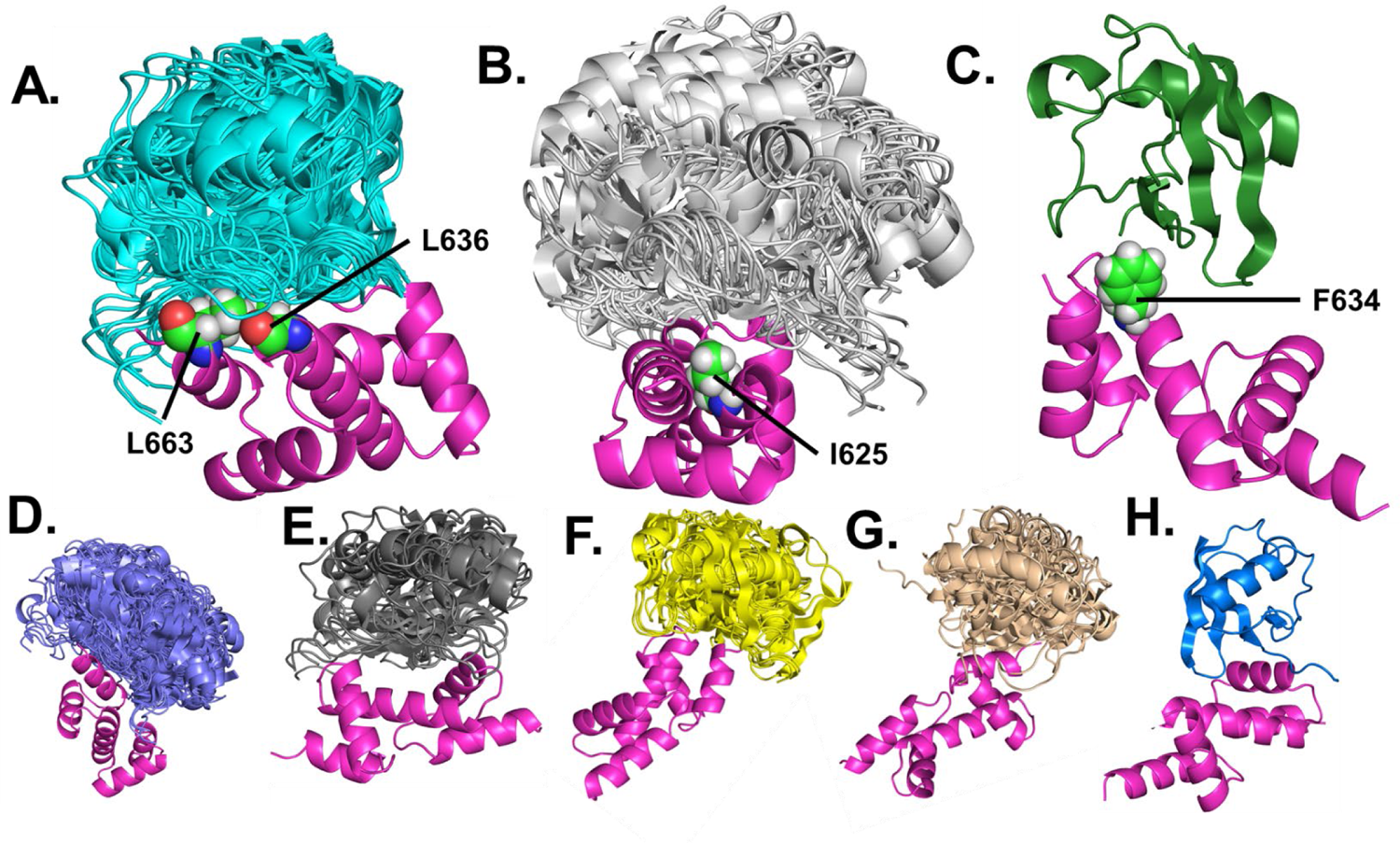
Computationally predicted UBA:ubiquitin binding surfaces. **A-C.** Binding site clusters that agreed with pull-down assays, involving UBA residues L636/L663 (A), I625 (B) and F634 (C). **D-H.** Clusters that were rejected because 1. mutagenesis of key amino acids showed no effect on ubiquitin binding, 2. they did not involve the canonical binding amino acids of ubiquitin (L8, I44, H68, and V70), 3. or unresolvable clashes were observed between the ubiquitin and the UBA domain.

**Figure S4:**
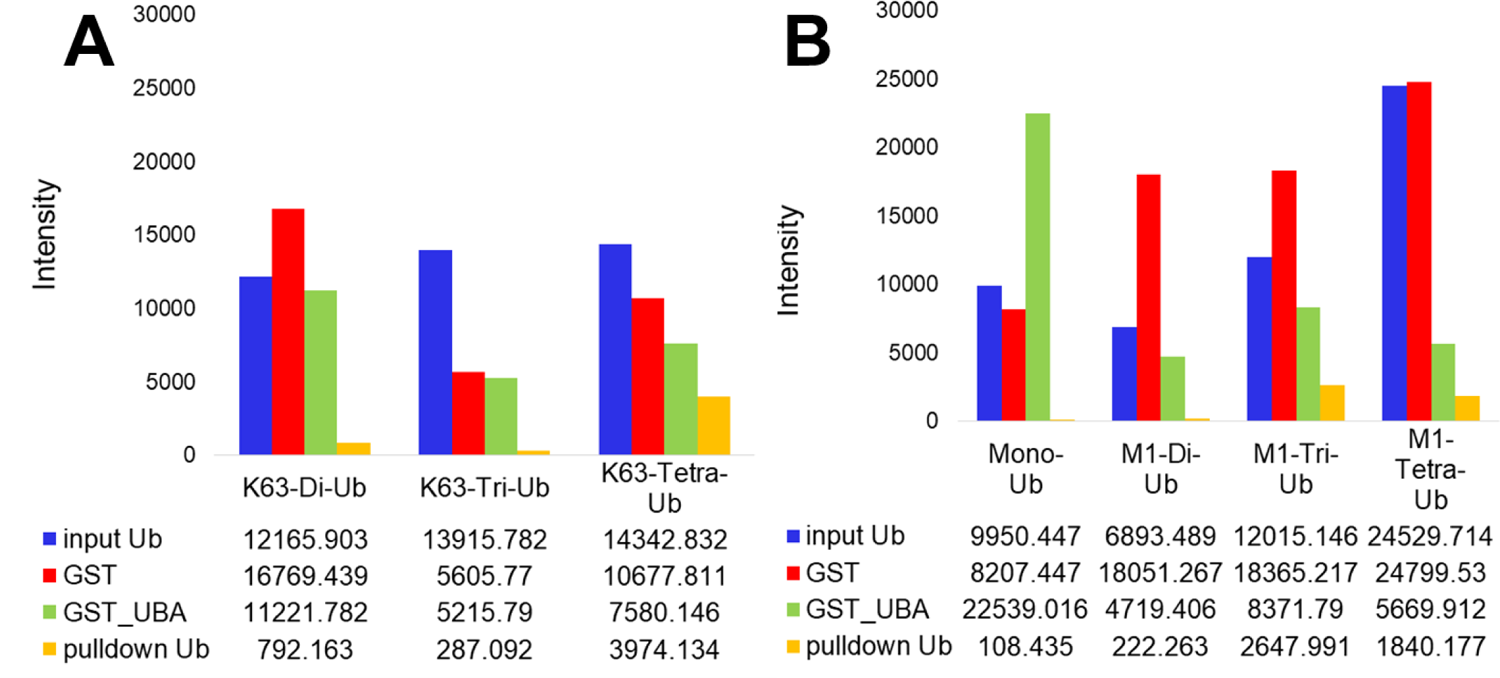
Quantification of the pulldown western blots using GST_UBA and K63- and M1-linked-ubiquitins. The input ubiquitin is shown in blue, GST is shown in red, GST_UBA is shown in green and the pulldown ubiquitin is shown in orange.

